# Identification of claudin-3 as an entry factor for rat hepacivirus

**DOI:** 10.1101/2025.02.27.640490

**Authors:** Tomohisa Tanaka, Yasunori Akaike, Hirotake Kasai, Atsuya Yamashita, Yoshiharu Matsuura, Kohji Moriishi

## Abstract

Approximately 58 million people worldwide are believed to be infected with hepatitis C virus (HCV), a major causative agent of chronic liver diseases. *Hepacivirus ratti* strain rn-1, which was discovered from *Rattus norvegicus* (Norway rat) and designated Norway rat hepacivirus 1 (NRHV1), shares similar properties with HCV in terms of genetic homology, target cell tropism, pathogenicity and the immune response. *In vivo* infection systems for NRHV1 will help overcome the challenges in HCV research for vaccine development. However, the virological characteristics of NRHV1, such as the mechanisms of cell entry, remain largely unexplored, in part owing to a paucity of cell culture systems for NRHV1. Here, we identified the host factors that facilitate NRHV1 entry by profiling the gene expression of two cell lines with different susceptibilities to NRHV1 infection. NRHV1 employs rodent orthologues of HCV entry factors, including scavenger receptor class BI, CD81, and occludin, and utilizes claudin-3 (CLDN3) but not claudin-1. The expression of rat and mouse, but not human, CLDN3 facilitates the entry of NRHV1 into murine cell lines that are insusceptible to NRHV1 infection. The host-specific cell entry of CLDN3 is determined by two amino acid residues, Ile^44^ and Trp^46^, in extracellular loop 1. These findings suggest that some hepacivirus species share an HCV-like entry mechanism but show evidence of adaptive evolution of their virus-specific entry factors to host animals.

## Introduction

Hepatitis C virus (HCV), a major causative agent of severe human liver diseases, is an enveloped positive-sense single-stranded RNA virus that belongs to the genus *Hepacivirus* of the family *Flaviviridae*. Approximately 58 million people are chronically infected with HCV, with an estimated 1.5 million new infections occurring each year ^1, 2^. The development of highly effective direct-acting antivirals (DAAs) has enabled the clearance of HCV in more than 95% of patients, but eliminating HCV worldwide is still challenging because of the high cost of DAAs and the absence of an effective vaccine. The development of the HCV replicon and infectious cell culture systems has substantially facilitated research on the life cycle of HCV and antiviral drugs ^3^. However, the paucity of versatile small animal models for HCV infection is a major obstacle to vaccine development and elucidation of the pathogenesis ^4^.

The genus *Hepacivirus* comprises more than 14 species, each of which has a limited host range, although some species have been reported to be transmitted between different hosts. Nonprimate hepacivirus (NPHV) was initially identified in dogs, but NPHV has since been predominantly detected in horses and can be naturally transmitted to donkeys ^5, 6, 7^. *Hepacivirus ratti* strain rn-1 was discovered in Norway rats (*Rattus norvegicus*) in New York; it was designated Norway rat hepacivirus 1 (NRHV1) and can also be transmitted to mice (*Mus musculus*) ^8, 9, 10^. NRHV1 and NPHV possess binding sites for liver-specific microRNA-122 (miR-122) in their 5’-untranslated regions (UTRs), similar to those in the HCV 5’UTR. The binding of miR-122 increases their replication, although stem loop I of the 5’UTR may support replication instead of miR-122 in the case of NPHV ^8, 11, 12, 13^. Furthermore, NRHV1 can persistently infect laboratory rat strains and immunocompromised murine lines, while the virus is eliminated four or five weeks after infection in immunocompetent mice. ^8, 10^. Transient injection of an antibody to CD4 before infection or continuous injections of an antibody to CD8 before and after infection have been shown to result in persistent NRHV1 infection in immunocompetent mice ^8, 14, 15^, suggesting that effective CD4 T-cell-dependent CD8 T-cell stimulation and the development of virus-specific antibodies are needed for NRHV1 clearance, which is similar to the immune response for spontaneous clearance of HCV in humans ^16, 17, 18^. Thus, a rat or murine infection system using NRHV1 is a useful surrogate model for HCV infection.

The entry of HCV into hepatocytes is known to be achieved through a multistep process. Initially, HCV attaches to the cell surface through interactions with proteoglycans, low-density lipoprotein receptor, scavenger receptor BI (SR-BI) and CD81. The virion is then translocated close to tight junctions via epithelial growth factor receptor signalling and internalized into cells, where it cooperates with occludin (OCLN) and claudin-1 (CLDN1) ^19, 20, 21^. *In vitro* studies have shown that CLDN6 and CLDN9 can support HCV entry instead of CLDN1 ^22, 23^. Several reports have suggested that SR-BI is also essential for NRHV1 entry and that GB virus-B (GBV-B) utilizes CLDN1 for entry ^24, 25, 26^. However, the other requirements in the entry step of hepaciviruses are still unclear. Recently, cell clones stably harbouring subgenomic replicon (SGR) RNA of NRHV1 were established from the rat hepatoma cell line McA-RH7777 and the mouse hepatoma cell line Hepa1-6 ^11^. The cured cells of SGR-harbouring McA-RH7777, McA-RH7777.hi, have been shown to permit infection with NRHV1 containing cell-adaptive mutations found in the replicon cell line ^26^, whereas other cell culture models for NRHV1 infection are still limited. Therefore, the NRHV1 life cycle and host–virus interactions have not yet been fully elucidated.

Here, we show that the cell line supporting the entire life cycle of NRHV1 is derived from the McA-RH7777 cell line but not from the murine hepatoma cell line AML12. Furthermore, our data suggest that NRHV1 basically utilizes CLDN3, but not CLDN1, for entry in addition to the rodent orthologues of the HCV entry factors SR-BI, CD81 and OCLN. The AML12 cell line permits NRHV1 entry following the exogenous expression of host CLDN3. Furthermore, our findings show that two adjacent amino acid residues that form the molecular surface of the first extracellular loop (EL1) of CLDN3 are essential for NRHV1 entry. These data provide insights into the mechanism of hepacivirus entry and suggest the evolutionary process by which hepaciviruses adapt to their individual hosts.

## Materials and Methods

### Cell lines

The McA-RH7777, AML12 and Hepa1-6 cell lines were maintained in Dulbecco’s modified Eagle’s medium (DMEM) supplemented with 10% foetal bovine serum (FBS, Gibco, Cambrex, MD, USA), 0.1 mM sodium pyruvate (Sigma–Aldrich, St. Louis, MO, USA), nonessential amino acids (Sigma–Aldrich) and 1% penicillin– streptomycin solutions (Nacalai Tesque, Kyoto, Japan). Plat-GP retrovirus packaging cells were maintained in medium supplemented with 10 µg/ml blasticidin (InvivoGen, San Diego, CA). The cell line harbouring NRHV1 SGR RNA was maintained in the presence of 500 µg/ml G418 (Nacalai Tesque). The cell lines stably expressing mCherry-MAVS and CLDN3 were selected with 500 µg/ml zeocin (InvivoGen) and 1 µg/ml puromycin (Nacalai Tesque), respectively. All the cells were cultured at 37°C in a 5% CO_2_ humidified atmosphere.

### Reagents and antibodies

Daclatasvir (DCV), pibrentasvir (PIB) and sofosbuvir (SOF) were purchased from Selleck (Houston, TX, USA) and dissolved in dimethyl sulfoxide (DMSO) for storage at -80°C until use. Rabbit polyclonal antibodies against the NRHV1 core and NS5A were commercially developed by immunization with the synthetic peptides SAAYPVRDPRRK and YETSNDHVPKEDSW, respectively (Scrum, Japan). Anti-CLDN1 (Invitrogen, RF217968), anti-CLDN3 (Abcam, ab15102), anti-SR-BI (Proteintech, 21277-1-AP), anti-OCLN (Abcam, ab216327), anti-β-actin (Sigma‒Aldrich, A5441), PE-conjugated anti-mouse/rat CD81 (BD Pharmingen, 559519), and PE-conjugated hamster IgG (BD Pharmingen, 553972) were purchased.

### Plasmids and transfection

The plasmid encoding the infectious clone of the NRHV1 rn1 strain, pRHV-rn1 ^10^, was kindly provided by Dr. Kapoor. The plasmid encoding the NRHV1 subgenomic reporter replicon (SGR), RHV-SGR-adFEO-10 ^11^, was kindly provided by Dr. Scheel. The lentiviral vector encoding miR-122 was prepared previously ^12, 27^. For the preparation of expression vectors, the ORFs of host factors were amplified by PCR using cDNA prepared from mouse liver and McA-RH7777 and Huh7 cells. The PCR amplicons were subsequently cloned and inserted into the BamHI and SalI sites of pEF pGKpuro ^28^ or the AflII and BamHI sites of pcDNA^TM^ 4 (Thermo Fisher Scientific, Waltham, MA). Site-directed mutagenesis was carried out via a KOD plus mutagenesis kit (Toyobo, Japan). The ORFs of rat CD81 and OCLN were inserted into the BamHI and EcoRI sites of the pMXs-IRES-GFP retroviral vector. For the specific knockouts of host factors, oligonucleotides containing sgRNA target sequences (Supplementary Table S1) were synthesized and cloned and inserted into the BbsI site of pSpCas9 (BB)-2A-Puro (Addgene #48139). Inserted sequences were confirmed by a DNA sequencing service (Fasmac, Japan). The expression vectors were transfected via Lipofectamine 3000 (Thermo Fisher Scientific).

### Gene knockdown

Predesigned siRNAs of the MISSION^TM^ were purchased from Merck and are listed in Supplementary Table S2. The universal negative control siRNA was purchased from Sigma-Aldrich. The cultured cells were transfected with 10 nM siRNAs using Lipofectamine RNAiMAX transfection reagent (Thermo Fisher Scientific).

### Preparation of virus

For production of infectious NRHV1, pRHV-rn1 ^10^ was linearized with MluI and transcribed *in vitro* via a MEGAscript T7 transcription kit (Thermo Fisher Scientific). The resulting RNA was introduced into Sprague‒Dawley rats via direct intrahepatic injection. Sera collected from the rats were intravenously injected into NOD-SCID mice. Sera containing NRHV1 (NRHV1ser) were collected from persistently infected mice, aliquoted and stored at -80°C. All animal experiments were approved by the Institutional Committee of Laboratory Animal Experimentation (University of Yamanashi) and performed in accordance with the guidelines for the care and use of laboratory animals (University of Yamanashi). The culture supernatant containing cultured NRHV1 (NRHV1cc) cells was filtered through a 0.45 μm filter, aliquoted and stored at -80°C. The amounts of NRHV1 RNA in the serum and culture supernatant were measured via qRT‒PCR as described below to estimate the viral load. Retrovirus-based pseudoparticles were produced in Plat-GP packaging cells via the cotransfection of a VSVG expression vector (pCMV-VSV-G) and pMXs vectors. The culture supernatants were harvested at 3 days post-transfection, filtered through a 0.45 μm filter, and stored at -80°C.

### Electroporation of replicon RNA, luciferase assay and colony formation assay

The plasmids encoding NRHV1-SGR-FEO with an adaptive mutation in NS5A (W503R) and with a replication-dead mutation in NS5B (GNN) ^11^ were linearized with MluI and transcribed *in vitro* via a MEGAscript T7 transcription kit (Thermo Fisher Scientific). The cultured cells (McA-RH7777, AML12 and Hep1-6) were removed by trypsinization. The removed cells were washed twice with ice-cold PBS. McA-RH7777 cells were resuspended in PBS at 1.5×10^7^ cells/ml, and 400 µl of cell suspension mixed with 6 µg of RNA was transferred into a 2 mm electroporation cuvette (Bio-Rad) and subjected to electroporation under the conditions described by Wolfisberg et al. ^11^. AML12 or Hepa1-6 cells were resuspended in PBS at 1×10^7^ cells/ml. Then, 800 µl of the cell suspension was mixed with 8 µg of RNA, transferred to a 4-mm electroporation cuvette (Cell Projects, Harrietsham, UK) and electroporated at 260 V and 975 µF using a Gene Pulser Xcell system (Bio-Rad). The electroporated cells were cultured as described above and collected at the indicated times, as described in the figure legends. For measurement of firefly luciferase activity in the transfected cells, the cells were washed once with PBS, harvested and lysed by freeze-thawing once in passive lysis buffer. Luciferase activity was measured using a Luminescencer-Octa (Atto, Japan) or a Glomax luminometer system (Promega). For the colony formation assay, the cells were cultured in the presence of G418 one day after electroporation. Colonies were fixed with cold methanol and stained with crystal violet.

### Quantification of viral and SGR RNAs and mRNAs

Total RNA was extracted from cells via an RNeasy Mini Kit (Qiagen, Hilden, Germany). Viral RNA in the supernatant was extracted via a QIAamp Viral RNA Mini Kit (Qiagen). The RNA samples were subsequently reverse transcribed to cDNA via a ReverTra Ace qPCR RT kit (Toyobo, Osaka, Japan). Quantitative PCR was carried out using Fast SYBR Green Master Mix (Applied Biosystems, Waltham, MA) and a QuantStudio 3 Real-Time PCR System (Applied Biosystems). The primers used are listed in Supplementary Table S3. The levels of viral RNAs and target mRNAs relative to GAPDH mRNA were calculated as delta Ct [ΔCt = Ct (target) − Ct (GAPDH)] and normalized to the level of the control group [ΔΔCt = ΔCt (experimental group) − ΔCt (control group)]. The relative levels were determined via the 2^−ΔCt^ or 2^−ΔΔCt^ method. The amount of NRHV1 RNA in the supernatant was quantified by developing a standard curve using known copy numbers of *in vitro*-transcribed NRHV1 RNA.

### Western blotting

The cells of interest were lysed on ice in lysis buffer (25 mM Tris-HCl pH 7.4, 150 mM NaCl, 1 mM EDTA, 0.5% SDS, 1% NP-40, and 5% glycerol) supplemented with a complete inhibitor cocktail (Roche). The lysate was mixed with one-third volume of 3× sample buffer, boiled and then subjected to SDS‒PAGE. The proteins in the gel were transferred onto a PVDF membrane (Millipore, Billerica, MA) using a semidry transfer blotter (ATTO, Tokyo, Japan). The membrane was incubated with Blocking One (Nacalai Tesque), reacted with appropriate antibodies and washed with TBS containing 0.02% Tween-20. The immune complexes were visualized with SuperSignal West Femto substrate (Pierce) and detected with an LAS-4000 Mini (GE Healthcare, Tokyo, Japan).

### Immunocytochemistry

The cells of interest were fixed on glass slides with 4% paraformaldehyde (PFA) at room temperature for 15 min, washed twice with phosphate-buffered saline (PBS) and incubated with PBS containing 5% FBS and 0.1% saponin at room temperature for 1 h. The resulting cells were then incubated with primary antibodies in dilution buffer (PBS containing 1% bovine serum albumin and 0.1% saponin). After being washed three times with PBS, the cells were incubated with secondary antibodies and DAPI in dilution buffer at room temperature for 1 h and washed three times with PBS. The fluorescence images were obtained by a fluorescence microscope (Keyence, Tokyo, Japan) or FluoView FV1000 laser confocal microscope (Olympus, Tokyo, Japan).

### Flow cytometry

The surface expression level of CD81 was measured by flow cytometry. The cells of interest were detached with trypsin-EDTA solution (Nacalai Tesque), suspended in culture medium, and then washed once with PBS. The resulting cells were incubated with PE-conjugated anti-CD81 or isotype IgG in FACS buffer (PBS containing 2% FBS) at 4°C for 1 h. Then, the resulting cells were washed twice with FACS buffer and analysed with a BD FACSCalibur. The data were analysed with CellQuest software.

### AlphaFold2 prediction and phylogenetic analysis

The amino acid sequences of CLDN3 and the mutant were submitted to ColabFold v1.5.5, running a simplified version of AlphaFold2 (https://colab.research.google.com/github/sokrypton/ColabFold/blob/main/AlphaFold2.ipynb) ^29, 30^. The rank 1 models were visualized using PyMOL v2.5.8 (Schrödinger, L. & DeLano, W., 2020. PyMOL, available at http://www.pymol.org/pymol). Phylogenetic analysis was conducted using MEGA11 ^31^. The sequences of the CLDN members were aligned via the MUSCLE algorithm. A phylogenetic tree was constructed via the neighbour‒joining method with 1000 bootstrap replicates.

### Statistics

The data used for the statistical analyses are expressed as the means ± standard deviations. The significance of differences between two groups was determined by Student’s t test. Significant differences among multiple groups were estimated by the Tukey‒Kramer test. A *p* value of less than 0.05 was considered statistically significant (*, *p* < 0.05; **, *p* < 0.01).

## Results

### Efficacy of NRHV1 replication in rodent cell lines

For determination of whether rodent hepatoma cell lines support NRHV1 replication, several cell lines were electroporated with adaptive NRHV1 SGR RNA encoding a fusion gene of firefly luciferase and neomycin phosphotransferase II, adFEO-10 ^11^, or nonreplicative SGR RNA with a GNN mutation in the NS5B coding region. The replication level was evaluated by transient and colony formation assays. NRHV1 SGR exhibited greater luciferase activity than did the GNN mutant in rat McA-RH7777 and murine Hepa1-6 cells (Fig. 1A). NRHV1 SGR also yielded G418-resistant colonies in these cell lines (Fig. 1B). These findings were consistent with the data described by Wolfisberg et al. ^11^, although the luciferase activity evaluated in the NRHV1 SGR-transfected AML12 cells was similar to that in the NRHV1 SGR GNN-transfected cells (Fig. 1A). Furthermore, AML12 cells electroporated with NRHV1 SGR were eradicated by G418 treatment (Fig. 1B). The efficacy of colony formation is shown in Table 1. These findings suggest that AML12 cells are incapable of supporting NRHV1 replication.

**Figure 1.**
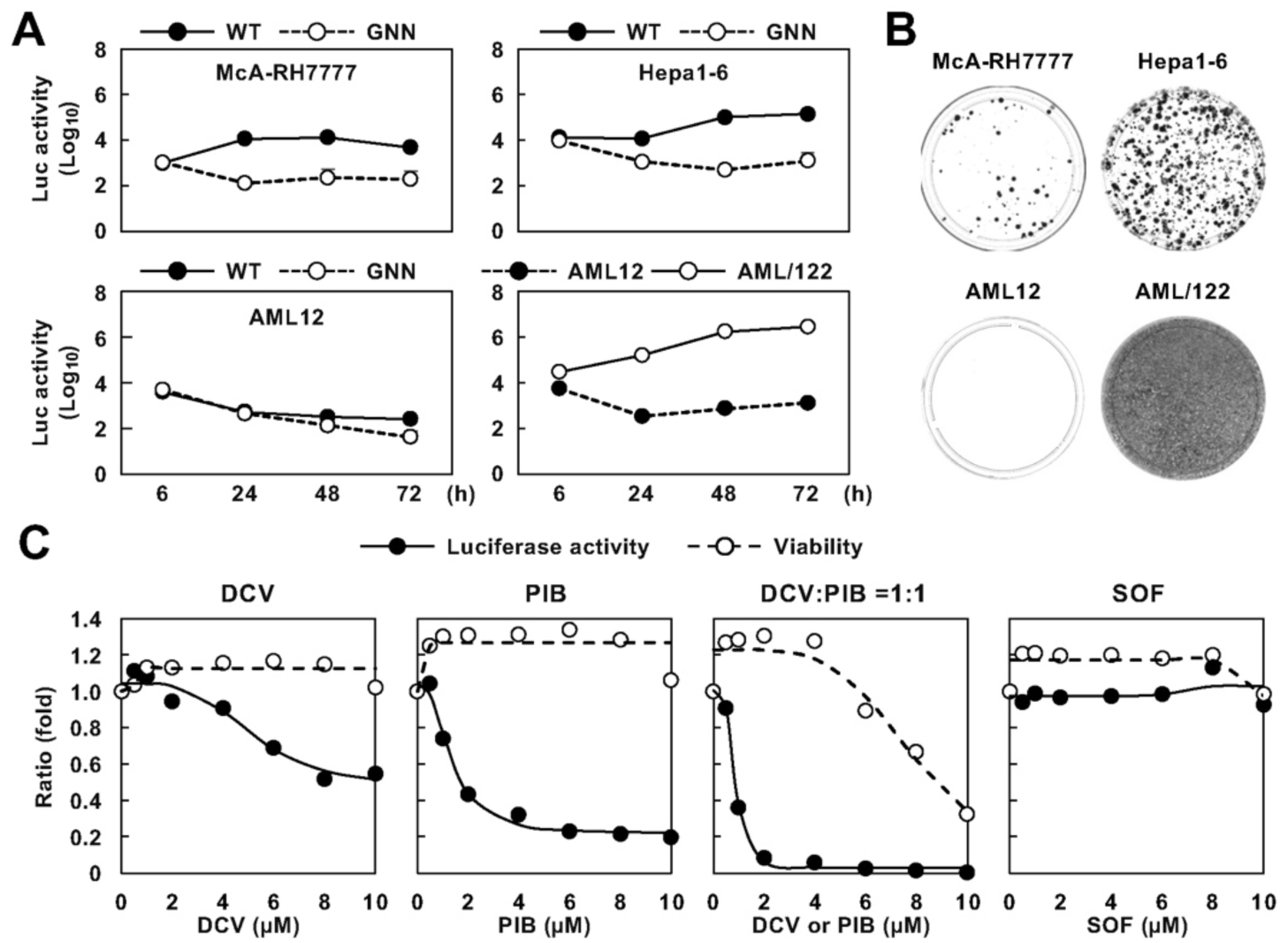
NRHV1 replication in rodent cells and antiviral agents against NRHV1. (A) Transient replication of the NRHV1 SGR in McA-RH7777, Hepa1-6, AML12 and AML/122 cells. Luciferase activities were measured at the indicated time points following electroporation with NRHV1 SGR (WT) or the nonreplicative replicon (GNN) RNA. (B) Colony formation assay of NRHV1 SGR in rodent cell lines. The cultured cells were electroporated with NRHV1 SGR, followed by selection with G418. G418-resistant colonies were stained with crystal violet. (C) McA-RH7777 cells harbouring NRHV1 SGR (McA/SGR#1) cells were incubated with DCV, PIB, SOF or a combination of DCV and PIB (DCV:PIB = 1:1, molar ratio) for 6 days at the indicated concentrations. The levels of replication and viability were determined by luciferase and MTS assays, respectively, and are shown as the values relative to those of the vehicle-treated groups.

**Table 1.** Colony formation unit (CFU) of cells transfected with NRHV1 SGR RNA.

| Cell | Host | CFU/μg RNA |
| --- | --- | --- |
| McA-RH7777 | Rat | $0.2 \times 10^3$ |
| Hepa1-6 | Mouse | $1.4 \times 10^3$ |
| AML12 | Mouse | 0 (< 10) |
| AML12/122 | Mouse | $5 \times 10^4$ |

Liver-specific miR-122 is a well-known host factor responsible for HCV replication, since exogenous expression of miR-122 can allow HCV replication in some cell lines that do not allow HCV replication ^32, 33, 34^. Given that NRHV1 replication is also dependent on miR-122 ^11^, we transduced AML12 cells with a lentiviral vector expressing miR-122 (AML/122). The expression level of miR-122 in the AML/122 cells was almost comparable to that in the mouse liver, whereas miR-122 was undetectable in the parental cells (Supplementary Fig. 1A). Transfection of NRHV1 SGR into AML/122 cells induced luciferase activity and yielded numerous G418-resistant colonies (Fig. 1A, B), suggesting that the nonpermissiveness of AML12 is due to miR-122 deficiency. We isolated two NRHV1 SGR-harbouring cell clones from each of the McA-RH7777, Hepa1-6 and AML/122 cell lines (Supplementary Fig. 1B, C).

We next treated McA-RH7777 cells harbouring NRHV1 SGR (McA/SGR) with several anti-HCV drugs and assessed the inhibitory effects on NRHV1 replication by a luciferase assay (Fig. 1C). The second-generation HCV NS5A inhibitor PIB inhibited replication in a dose-dependent manner (50% inhibitory concentration [IC_50_] = 1.7 μM), whereas the NS5B inhibitor SOF was not effective at a concentration of 10 μM, which was consistent with previous observations ^11^. In this study, we found that the first-generation NS5A inhibitor DCV also inhibited replication (IC_50_ > 10 μM). A 1:1 mixture of DCV and PIB was added to the supernatant. Compared with single treatment, treatment with the mixture was more effective without apparent cell toxicity (each IC_50_ = 0.9 μM and each IC_90_ = 1.6 μM). Therefore, the combination treatment was employed to establish the cured cell line. Several surviving clones were established from each SGR-harbouring cell line via treatment with 2 μM DCV or PIB for 2 weeks. NRHV1 SGR was not detected in these clones by a luciferase assay or RT‒qPCR evaluation (Supplementary Fig. 1B, C). In addition, these cured clones were susceptible to G418 treatment (data not shown). We determined that these surviving clones lacked the replicon and were identified as cured cells in this study.

### Permissibility of rodent cell clones for NRHV1 infection

Given that the replication efficacy of AML/122 cells was greater than that of murine Hepa1-6 cells (Fig. 1A, B), we investigated whether the cured cell clones derived from McA-RH7777- or AML/122-based replicon cells would permit NRHV1 infection. The cell clones were inoculated with the sera of the mice infected with NRHV1 (NRHV1ser) and then passaged every 3 or 4 days. The cells and supernatants were collected during serial passaging, and NRHV1 RNA was measured via qRT‒PCR. The level of intracellular viral RNA gradually increased at 31 days post-inoculation (dpi) in the McA1.8 cell clone but not in the McA1.5, McA4.2 or McA4.4 cell clones (Fig. 2A). Accordingly, the amount of NRHV1 RNA in the supernatant increased in a time-dependent manner in the McA1.8 cells (Fig. 2B). NS5A-positive foci were found in NRHV1ser-inoculated McA1.8 cells but not in naïve cells (Fig. 2C). The level of intracellular NRHV1 RNA was reduced by treatment with PIB, suggesting active replication of the genomic viral RNA in McA1.8 cells after entry (Fig. 2D). We next determined whether cell culture-derived infectious NRHV1 (NRHV1cc) was released from the cells. The culture supernatant of McA1.8 cells infected with NRHV1ser at 35 dpi was inoculated into naïve McA1.8 cells. NS5A-positive cells were subsequently observed after inoculation (Fig. 2E), whereas NS5A and core were detected in the inoculated cells by Western blotting (Fig. 2F). Furthermore, the amount of intracellular and supernatant NRHV1 RNA logarithmically increased in inoculated McA1.8 cells in a time-dependent manner (Fig. 2G, H). These data suggest that the NRHV1 virus is included in the culture supernatant of the inoculated cells. Serial passages of NRHV1ser in McA1.8 cells improved viral fitness *in vitro*. We determined the differences in the nucleotide sequences of NRHV1ser and NRHV1cc and listed the nonsynonymous mutations observed in the NRHV genome in Supplementary Table S4 (Reference genome: KX905133).

**Figure 2.**
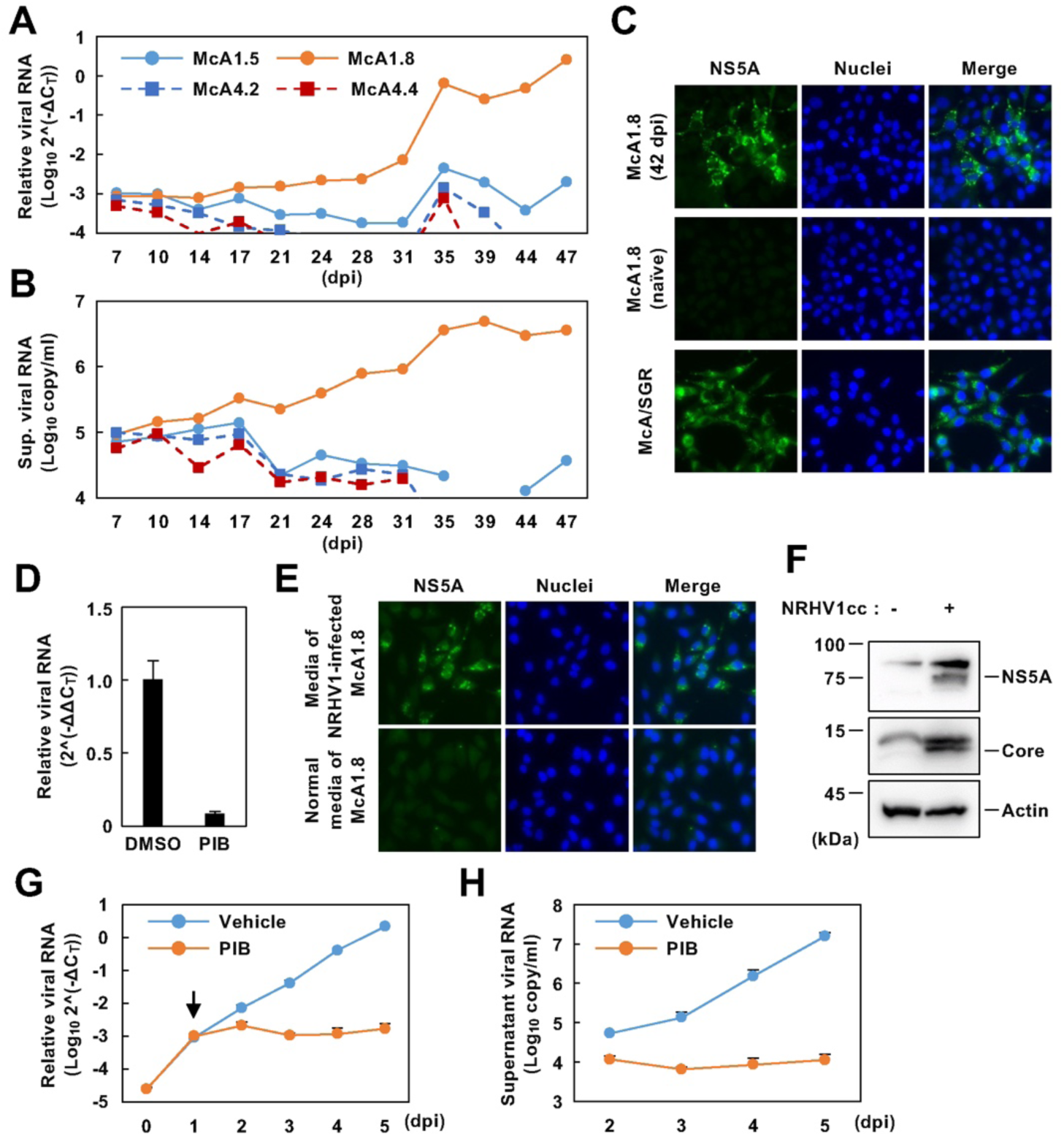
Establishment of a rat cell line that produces infectious particles of NRHV1. (A, B) McA-RH7777-derived cured cell clones were inoculated with the NRHV1ser at 6 copies/cell and were passaged every three or four days. The intracellular NRHV1 RNA level relative to the GAPDH mRNA level (ΔCt) (A) and the amount of NRHV1 RNA in the supernatant (B) were quantified via qRT‒PCR. (C) NRHV1ser-inoculated McA1.8 cells (42 dpi), naïve McA1.8 cells and McA/SGR#1 cells were fixed on glass chamber slides and stained with an anti-NS5A antibody and DAPI. (D) NRHV1ser-inoculated McA1.8 cells were incubated with 5 µM PIB from 31 to 35 dpi and subjected to qRT‒PCR. The NRHV1 RNA level relative to the GAPDH mRNA level (ΔCt) was normalized to that in cells treated with DMSO (ΔΔCt). (E) Culture media of NRHV1ser-inoculated McA1.8 cells were added to naïve McA1.8 cells at 5 copies/cell (upper panels). Normal medium from naïve McA1.8 cells was used as a negative control (lower panels). These cells were fixed at 7 dpi and stained with an anti-NS5A antibody and DAPI. (F) Naïve McA1.8 cells were incubated with normal media or media containing NRHV1cc (5 copies/cell). These cells were harvested at 4 dpi and subjected to Western blotting. (G, H) Time course study of NRHV1cc propagation. McA1.8 cells were inoculated with media containing NRHV1cc at 2.5 copies/cell. These cells were washed three times with fresh medium, and the medium was replaced with medium containing vehicle (DMSO) or 2 μM PIB at 1 dpi. The intracellular NRHV1 RNA level relative to the GAPDH mRNA level (ΔCt) (G) and the amount of supernatant NRHV1 RNA (H) were quantified via qRT‒PCR. The arrow in (G) indicates the point of PIB addition.

Cured cells, which were prepared from replicon cells harbouring HCV SGR following treatment with antivirals, can support HCV replication more effectively than can parental cells ^35^. McA1.8 and parental cells were electroporated with synthesized NRHV1 SGR. Compared with the parental cells, the McA1.8 cells presented greater luciferase activity at 24–48 h post-transfection (Supplementary Fig. 2A), while colony formation efficacy was also greater in the McA1.8 cells than in the parental cells (Supplementary Fig. 2B). The level of miR-122 was greater in the McA1.8 cells than in the parental cells (Supplementary Fig. 1A). These data suggest that the increased permissiveness for NRHV1 replication facilitated viral propagation in the McA1.8 cells in a similar way to that observed in the HCV study. In contrast, the level of intracellular NRHV1 RNA did not increase in any of the cured clones derived from AML/122 after two months (Supplementary Fig. 2C). As shown in Fig. 1A and B, AML/122 cells strongly supported NRHV1 replication. However, NRHV1 RNA was not detected even in the cured cell lines after inoculation with cell culture-adapted NRHV1cc prepared from McA1.8 cells (Supplementary Fig. 2D). These data suggest that AML12-derived clones lack sensitivity to NRHV1 entry or disassembly.

### Involvement of rodent orthologues of HCV entry factors in NRHV1 entry

SR-BI is required for NRHV1 infection ^26^, but other host factors involved in NRHV1 entry have not yet been identified. Given that CD81, OCLN, CLDN1 and SR-BI are well characterized as the main entry factors for HCV infection ^19, 21^, we investigated the roles of these rodent orthologues in NRHV1 entry. We initially evaluated the expression levels of rodent orthologues in NRHV1-permissible (McA1.8) and nonpermissible cell lines (Hepa1-6 ^11^ and AML/122). Total RNA extracted from murine livers was used as a positive control because mouse hepatocytes permit NRHV1 infection ^8^. The expression levels of SR-BI and CD81 were comparable among all the rodent cells used in the present study, whereas OCLN was expressed in McA1.8 cells but not in the murine cell lines (Fig. 3A). CLDN1 was detected in mouse livers but not in all cell lines (Fig. 3A), suggesting that CLDN1 is not necessary for NRHV1 entry. Next, we examined the role of SR-BI, CD81 and OCLN in NRHV1cc infection. SR-BI, CD81 or OCLN mRNA levels were significantly decreased at 24 h after transfection with the respective siRNAs (Fig. 3B). Compared with those in mock cells, the intracellular NRHV1 RNA levels were significantly lower at 3 dpi when NRHV1cc was inoculated into the cells 24 h after siRNA transfection (Fig. 3C). Transfection with si-CD81#2, which is siRNA targeting CD81 mRNA, did not affect NRHV1 RNA levels because of the low knockdown efficacy (Fig. 3B, C). SR-BI knockout reportedly impairs NRHV1 infection both *in vivo* and *in vitro* ^25, 26^, although whether CD81 knockout or OCLN knockout affects NRHV1 infection is not known. We next established CD81- or OCLN-knockout cell lines from McA1.8 cells (Supplementary Fig. 3A, B) to test the requirement of both proteins for NRHV1 infection. NRHV1 propagation was impaired, but not completely inhibited, in CD81-knockout cell clones (Fig. 3D). In contrast, OCLN-knockout cell lines did not show susceptibility to NRHV1cc infection (Fig. 3E). When CD81- and OCLN-knockdown clones were transduced with retroviral vectors encoding CD81 and OCLN, respectively, their susceptibility to NRHV1cc infection was restored (Fig. 3F, G and Supplementary Fig. 3C, D). The level of NRHV1 SGR in McA/SGR cells was not affected by any siRNA transfection (Supplementary Fig. 4A), suggesting that SR-BI, CD81 and OCLN are not involved in NRHV1 replication. These findings suggest that CD81, OCLN, and SR-BI, but not CLDN1, play important roles in NRHV1 infection.

**Figure 3.**
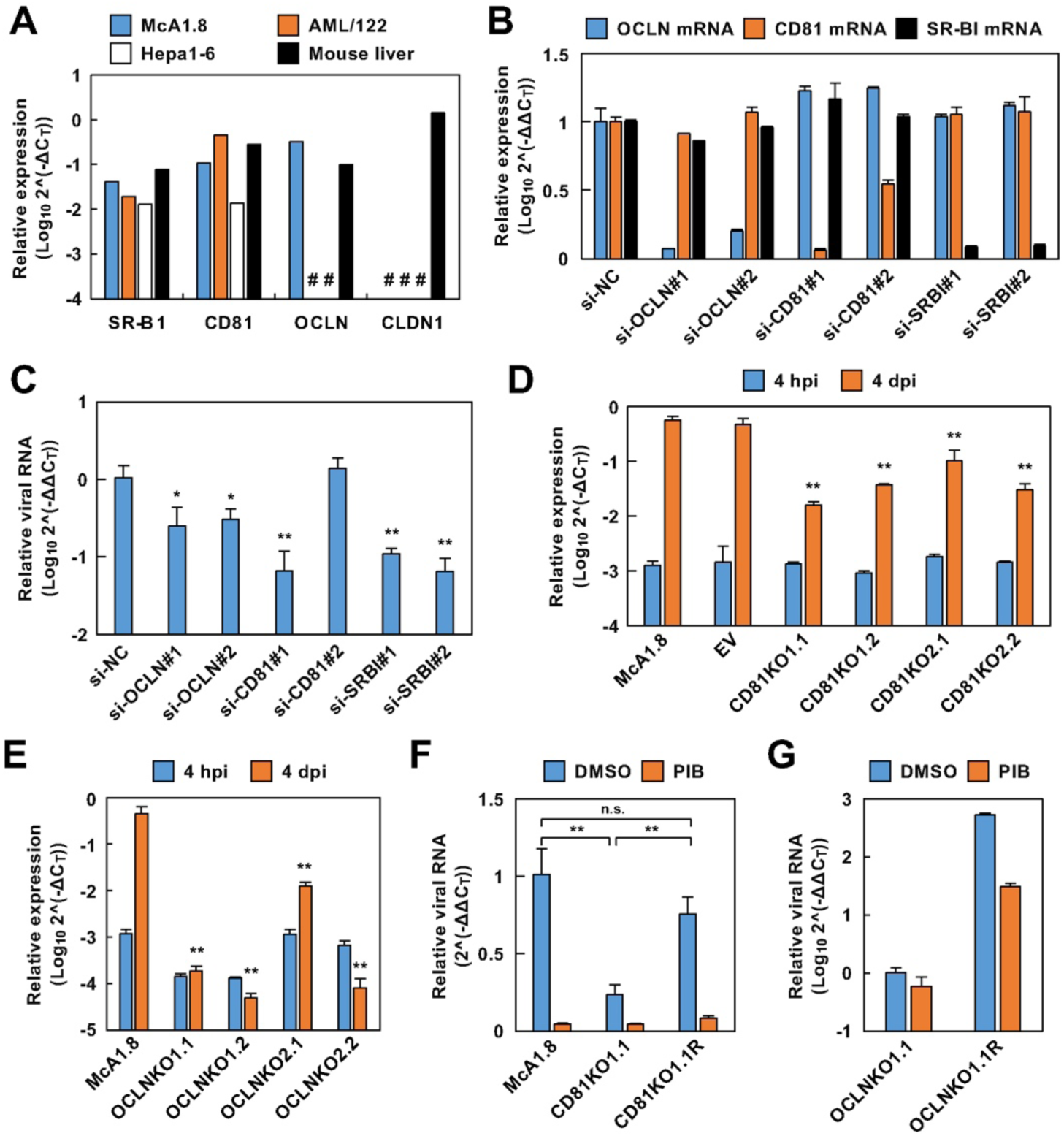
Rodent orthologues of HCV entry factors required for NRHV1 entry. (A) Total RNA extracted from the indicated cells and mouse livers was subjected to qRT‒ PCR to target the mRNAs of SR-BI, CD81, OCLD, and CLDN. The levels of mRNAs relative to those of GAPDH (ΔCt) were plotted. Undetectable values are indicated by number signs (#). (B, C) McA1.8 cells were transfected with 10 µM siRNA targeting OCLD, CD81 or SR-BI mRNA 24 h before inoculation with NRHV1cc at 5 copies/cell. The transfected cells were harvested at 0 dpi (B) or 3 dpi (C) and subjected to qRT‒ PCR. The levels of each mRNA or NRHV1 RNA relative to GAPDH mRNA (ΔCt) were normalized to the levels in cells transfected with the control siRNA, si-NC (ΔΔCt). The asterisks indicate significant differences in NRHV1 RNA compared with the control, as estimated by the Tukey‒Kramer multiple comparison test (*, *p* < 0.05. **, *p* < 0.01). (D, E) CD81-knockout (KO) or OCLN-KO cell lines were obtained from McA1.8 cells using two different sgRNAs for each gene. Empty vector-transfected cell lines (EVs) were used as controls. These cell lines were inoculated with NRHV1cc at 5 copies/cell and then harvested at 4 hpi or 4 dpi. Total RNA was subjected to qRT‒PCR. The asterisks indicate significant differences in NRHV1 RNA compared with that in McA1.8 cells at 4 dpi, as estimated by the Tukey‒Kramer multiple comparison test (**, *p* < 0.01). (F) CD81KO1.1 and rescued (CD81KO1.1R) cells were inoculated with NRHV1cc at 5 copies/cell and then cultured for 3 days in the presence or absence of 2 μM PIB. Significant differences between the indicated groups were estimated via the Tukey‒Kramer multiple comparison test (n.s., not significant). **, *p* < 0.01). (G) OCLNKO1.1 and rescued (OCLNKO1.1R) cells were inoculated with NRHV1cc at 5 copies/cell and then cultured for 3 days in the presence or absence of 2 μM PIB. (F, G) Total RNA was subjected to qRT‒PCR. The levels of NRHV1 RNA relative to those of GAPDH mRNA (ΔCt) were normalized to the levels in vehicle-treated KO cells (ΔΔCt).

### CLDN3 is involved in NRHV1 entry

CLDN1 is an entry factor for HCV infection of human hepatocytes, while CLDN6 and CLDN9 have been reported to be capable of supporting HCV entry *in vitro* ^22, 23^. Thus, we hypothesized that CLDN member(s) other than CLDN1 are involved in NRHV1 infection. We selected which CLDN family member is expressed in hepatocytes on the basis of single-cell RNA sequence data from mouse hepatocytes ^36^ and the NCBI database. The expression levels of these genes were surveyed in two rodent cell lines and mouse livers (Fig. 4A). The levels of CLDN2, CLDN6, CLDN12, CLDN15 and CLDN25 were comparable among the three groups. Rodent cell lines lacked the expression of CLDN5 and CLDN26. The levels of CLDN9, CLDN14 and CLDN24 were greater in murine cells and livers than in rat cells. Interestingly, CLDN3 and CLDN23 were expressed in NRHV1-permissible McA1.8 cells and mouse livers but not in nonpermissible AML/122 cells. McA1.8 cells were transfected with siRNAs targeting CLDNs prior to NRHV1cc infection (Fig. 4B). We silenced CLDN2 as a reference gene because of its comparable expression level between the two cell lines. The level of intracellular NRHV1 RNA in NRHV1cc-infected McA1.8 cells was specifically reduced by CLDN3 knockdown (Fig. 4C). However, the level of NRHV1 SGR RNA was not affected by CLDN knockdown (Supplementary Fig. 4B). The phylogenetic analysis of CLDN family members suggested that CLDN3 is highly related to CLDN6 and CLDN9 and relatively different from CLDN1 (Supplementary Fig. 4C). We generated CLDN3-knockout McA1.8 (C3KO) cell lines using CRISPR-Cas9 to examine the involvement of CLDN3 in NRHV1 entry (Fig. 4D and Supplementary Fig. 4D). C3KO cells were completely resistant to NRHV1cc infection (Fig. 4E), and the CLDN3 mutant resistant to sgRNA was capable of rescuing NRHV1cc infection in C3KO cells (Fig. 4F and Supplementary Fig. 4E). Taken together, these data suggest that CLDN3 mediates NRHV1 entry.

**Figure 4.**
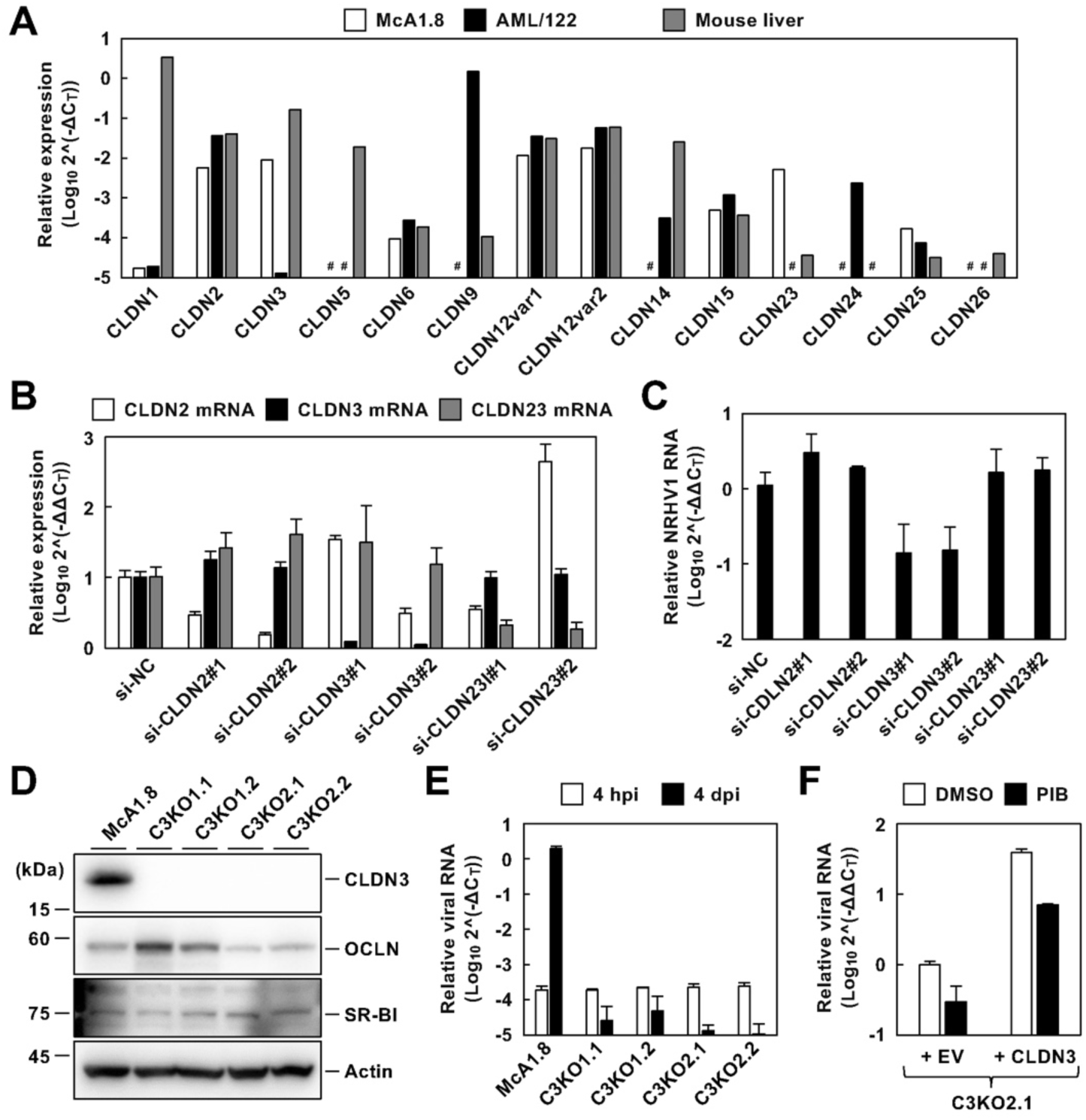
Identification of CLDN members that facilitate NRHV1 infection. (A) Total RNA extracted from the indicated cell lines and mouse livers was subjected to qRT‒PCR. The level of an individual claudin mRNA relative to that of GAPDH (ΔCt) was plotted. Undetectable values are indicated by number signs (#). (B) McA1.8 cells were transfected with 10 µM siRNAs targeting CLDN2, CLDN3 or CLDN23 mRNA 24 h before inoculation with NRHV1cc at 5 copies/cell. The transfected cells were harvested at 0 dpi (B) or 3 dpi (C) and subjected to qRT‒PCR. The levels of an individual mRNA or NRHV1 RNA relative to GAPDH mRNA (ΔCt) were normalized to the levels in cells transfected with si-NC (ΔΔCt). (D) Western blot analysis of lysates extracted from the indicated cells. (E) The indicated cells were inoculated with NRHV1cc at 5 copies/cell and then harvested at 4 hpi or 4 dpi. The levels of viral RNA were evaluated via qRT‒PCR. (F) C3KO cells were transfected with a plasmid encoding sgRNA-resistant CLDN3 24 h before inoculation with NRHV1cc at 5 copies/cell. These cells were cultured in the presence or absence of 2 µM PIB and harvested at 3 dpi. The levels of viral RNA were evaluated via qRT‒PCR.

### Establishment of NRHV1-susceptible murine cell lines

We hypothesized that NRHV1 does not infect AML/122 cells due to low or absent expression of CLDN3. We utilized the mitochondrial antiviral-signaling protein (MAVS)-based reporter system for HCV infection for real-time monitoring of NRHV1 infection ^37^. NRHV1 NS3/4A, but not the protease-inactive mutant harbouring the S138A mutation ^38^, cleaved mouse MAVS (Supplementary Fig. 5A), which is consistent with findings reported by Anggakusuma et al. ^39^. Then, we prepared McA1.8 or AML/122 cell clones stably expressing the fluorescent reporter protein composed of mCherry, the nuclear localization signal (NLS) and the C-terminal region of mouse MAVS (mCnls-MAVS) (Supplementary Fig. 5B). Although mCnls-MAVS is primarily localized on the outer membrane of mitochondria, the expression of NS3/4A, but not the S138A mutant, triggered the cleavage of mCnls-MAVS to release mCherry-NLS (mCnls), resulting in the translocation of mCnls into the nucleus (Supplementary Fig. 5C). Nuclear translocation of mCnls was observed in McA1.8-derived reporter cells following NRHV1cc infection (Supplementary Fig. 5D), suggesting that the fluorescent reporter system distinguishes NRHV1-infected cells from naïve cells. The intranuclear localization of mCnls was also detected in AML/122-derived reporter cells transiently expressing NS3/4A (Supplementary Fig. 5E). We next investigated whether the enforced expression of rat OCLN (rOCLN) and rat CLDN3 (rC3) confers NRHV1 susceptibility to AML/122 cells. AML/122 cells were cotransfected with OCLN and CLDN3 plasmids. The fluorescence was retained in the cytoplasm in the NRHV1cc-inoculated AML/122 reporter cells, whereas the nuclear translocation of mCnls was detected in the cells transiently expressing OCLN and CLDN3 (Supplementary Fig. 5F). AML/122 cells transfected with CLDN3, but not the OCLN vector, exhibited susceptibility to NRHV1 (Fig. 5A). Furthermore, AML/122 cell clones stably expressing CLDN3 (AML/122/rC3) exhibited susceptibility to NRHV1cc (Fig. 5B, C). The nuclear translocation of mCnls was observed in AML/122/rC3 cells expressing mCnls-MAVS after incubation with NRHV1cc (Fig. 5D). Serial passages of NRHV1cc-infected AML/122 cells demonstrated persistent NRHV1 replication, although NRHV1 RNA was undetectable in the supernatants (Supplementary Fig. 6A, B). We next determined whether infectious NRHV1cc particles were assembled in AML/122/rC3 cells. The cell extracts from NRHV1cc-infected AML/122/rC3 and McA1.8 cells were collected after freeze‒thawing cycles (Fig. 5E) and inoculated into naïve McA1.8 cells according to the methods of Gastaminza et al. ^40^. The amount of NRHV1 RNA in the cell extracts roughly correlated with the level of intracellular NRHV1 RNA (Fig. 5F). The extract from the McA1.8 cells was diluted to adjust the amount of NRHV1 RNA present in the cell extracts. The level of intracellular NRHV1 RNA increased in naïve AML/122/rC3 cells inoculated with the NRHV1-infected McA1.8 cell extract but not in those inoculated with the NRHV1cc-infected AML/122/rC3 cell extract (Fig. 5G). The nuclear translocation of mCnls was observed in the McA1.8/mCnls-MAVS cells inoculated with NRHV1cc-infected McA1.8 cell extract but not in those inoculated with the AML/122/rC3 cell extract (Fig. 5H), similar to the data shown in Fig. 5G. Thus, the replenishment of CLDN3 increases the susceptibility of AML/122 cells to NRHV1, while the process of viral particle assembly is suggested to be disrupted in murine cell lines. In contrast, the replenishment of OCLN promoted NRHV1cc infection in AML/122/rC3 cells but not in AML/122 cells (Fig. 5I), suggesting that there is an unknown factor for the entry function instead of OCLN in AML/122 cells.

**Figure 5.**
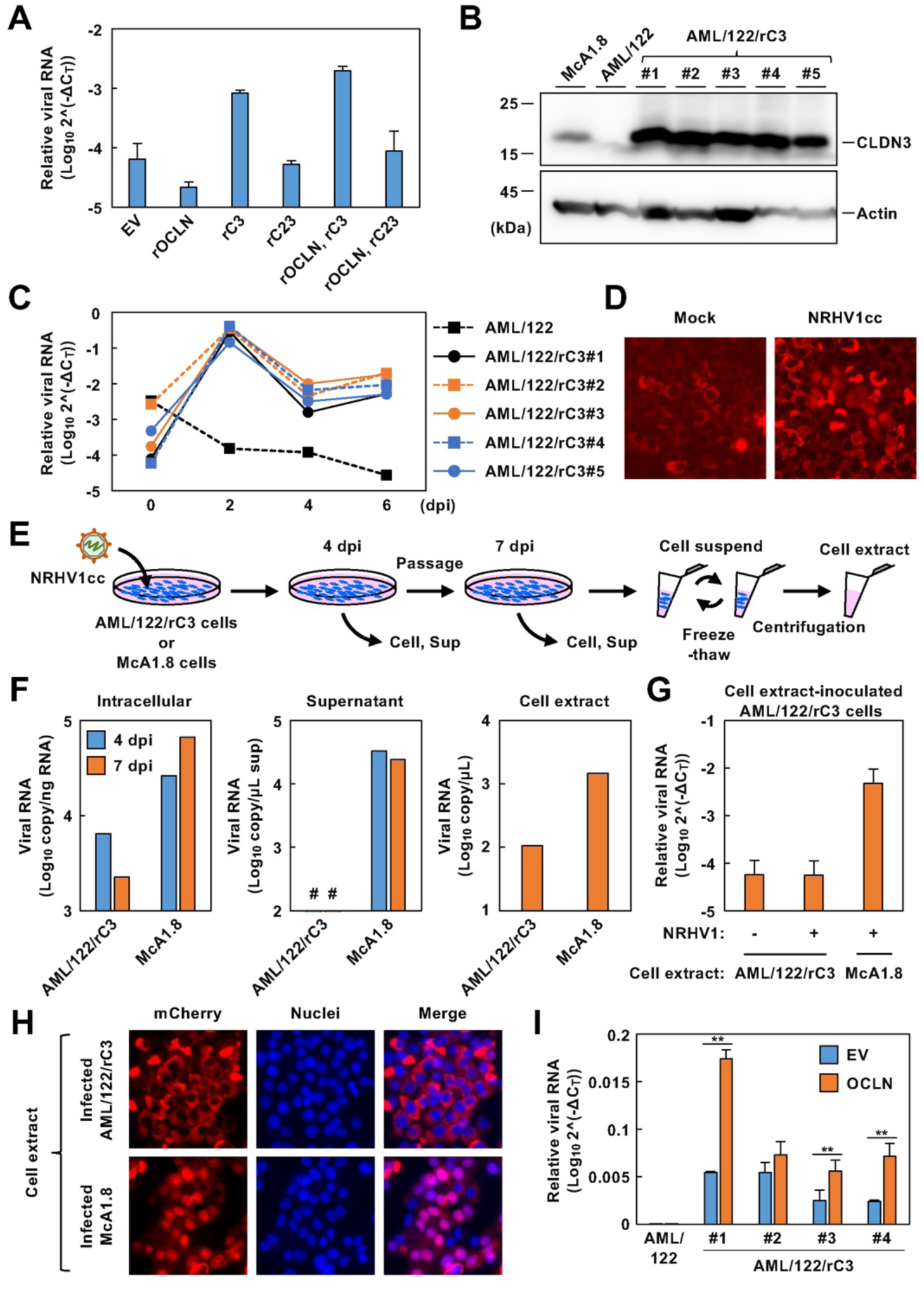
NRHV1 susceptibility of AML12-derived cells expressing CLDN3. (A) AML/122 cells were transfected with the indicated expression plasmid 24 h before inoculation with NRHV1cc (5 copies/cell). The transfected cells were harvested at 3 dpi, and the NRHV1 RNA level relative to the GAPDH mRNA level (ΔCt) was evaluated via qRT‒PCR. (B) Western blot analysis of lysates extracted from the indicated cell clones. The numbers indicate the number of cell clones. (C) These cell clones were incubated with NRHV1cc at 5 copies/cell for 5 h and washed with PBS to remove residual uninfected virus. The cells of an individual clone were harvested at the indicated time points. The intracellular NRHV1 RNA level relative to the GAPDH mRNA level (ΔCt) was quantified via qRT‒PCR. (D) AML/122/rC3 cells expressing mCnls-MAVS were infected with mock or NRHV1cc (10 copies/cell) and then fixed at 4 dpi. Images of these cells were obtained using a fluorescence microscope after staining. (E) Preparation of intracellular infectious particles. The cell extracts were isolated from NRHV1-infected AML/122/rC3 and McA1.8 cells at 7 dpi through freeze‒thaw cycles. The extracts were then inoculated into naïve AML/122/rC3 or McA1.8/mCnls-MAVS cells at 1 copy/cell. (F) NRHV1 RNA in cells, supernatants and cell extracts was evaluated via qRT‒PCR. (G) These extracts of an individual infected cell line (+) or mock (-) were inoculated into naïve AML/122/rC3 cells at 1 copy/cell and harvested at 2 dpi. NRHV1 RNA levels relative to those of GAPDH (ΔCt) were determined via qRT‒PCR. (H) Naïve McA1.8/mCnls-MAVS cells were inoculated with cell extracts at 1 copy/cell. These cells were fixed at 5 dpi and stained with DAPI. (I) An individual cell clone was transfected with a plasmid encoding rOCLN or an empty vector (EV) 24 h before inoculation with NRHV1cc at 5 copies/cell. These cells were harvested at 3 dpi. Total RNA was subjected to qRT‒PCR. * *p* < 0.05, ** *p* < 0.01 by Student’s t test.

### Species-specific and cross-species residues of CLDN3 that are crucial for NRHV1 entry

Since mouse, rat and hamster CLDN1, but not human CLDN3 (hCLDN3), could support HCV entry ^20, 23, 41, 42^, we investigated the specificity of CLDN3 as an entry factor for NRHV1. The transient expression of mouse CLDN3 (mCLDN3) and rat CLDN3 (rCLDN3), but not hCLDN3 or mCLDN1, conferred NRHV1 susceptibility to AML/122 cells (Fig. 6A, B). CLDN3 is composed of four transmembrane regions with two extracellular loops (EL1 and EL2) and a C-terminal tail domain (CTD) ^43^, and there are species-specific amino acid residues throughout the sequence (Supplementary Fig. 7A). We constructed three chimeric proteins between hCLDN3 and rCLDN3 (Fig. 6C) to identify the region responsible for entry. AML/122 cells were susceptible to NRHV1 when the hCLDN3 chimaera containing rat EL1 was expressed but not when the EL2 and CTD were expressed (Fig. 6D and Supplementary Fig. 7B), suggesting that the EL1 domain is responsible for species-specific receptor function. Given the presence of five distinct amino acid residues between the rCLDN3 and hCLDN3 EL1 domains, each amino acid residue within the hCLDN3 EL1 domain was substituted with its counterpart from rCLDN3 (Supplementary Fig. 7A). Only the N44I mutant of hCLDN3 conferred NRHV1 susceptibility to AML/122 cells (Fig. 6E and Supplementary Fig. 7C). In contrast, the I44N mutant of rCLDN3 presented a reciprocal phenotype (Fig. 6F and Supplementary Fig. 7D). These findings suggest that Ile^44^ in the EL1 domain contributes to the host-specific function of CLDN3 as an NRHV1 entry factor. AlphaFold2 prediction indicated that Ile^44^ constituted the second β-strand and was exposed on the surface (Supplementary Fig. 8A). The introduction of the I44N mutation (substitution of Ile^44^ with Asn) was predicted to result in the interaction between Asn^44^ and Phe^34^, which altered the surface structure of the EL1 domain (Supplementary Fig. 8B). Next, the residues proximal to Ile^44^ on the molecular surface were substituted with alanine (Supplementary Fig. 8C). The W46A mutant (substitution of Trp^46^ with Ala) did not confer susceptibility to NRHV1cc (Fig. 6G and Supplementary Fig. 8D). Trp^46^ is conserved across species and is predicted to be adjacent to Ile^44^ (Supplementary Figs. 7A, 8C). Thus, a species-dependent microstructure comprising Ile^44^ and Trp^46^ on CLDN3 is suggested to be essential for NRHV1 entry. As EL1 of CLDN1 is required for the internalization of the HCV E2-CD81 complex ^19, 44^, the interactions between NRHV1 E2 and CD81 and between CD81 and CLDN3 were analysed. As expected, CD81 and NRHV1 E2 were colocalized on the cell surface (Fig. 6H). Similarly, CD81 and CLDN3 also colocalized (Fig. 6I), suggesting that CLDN3 plays an important role in the recruitment of the NRHV1 E2-CD81 complex to the late-stage entry complex required for viral internalization.

**Figure 6.**
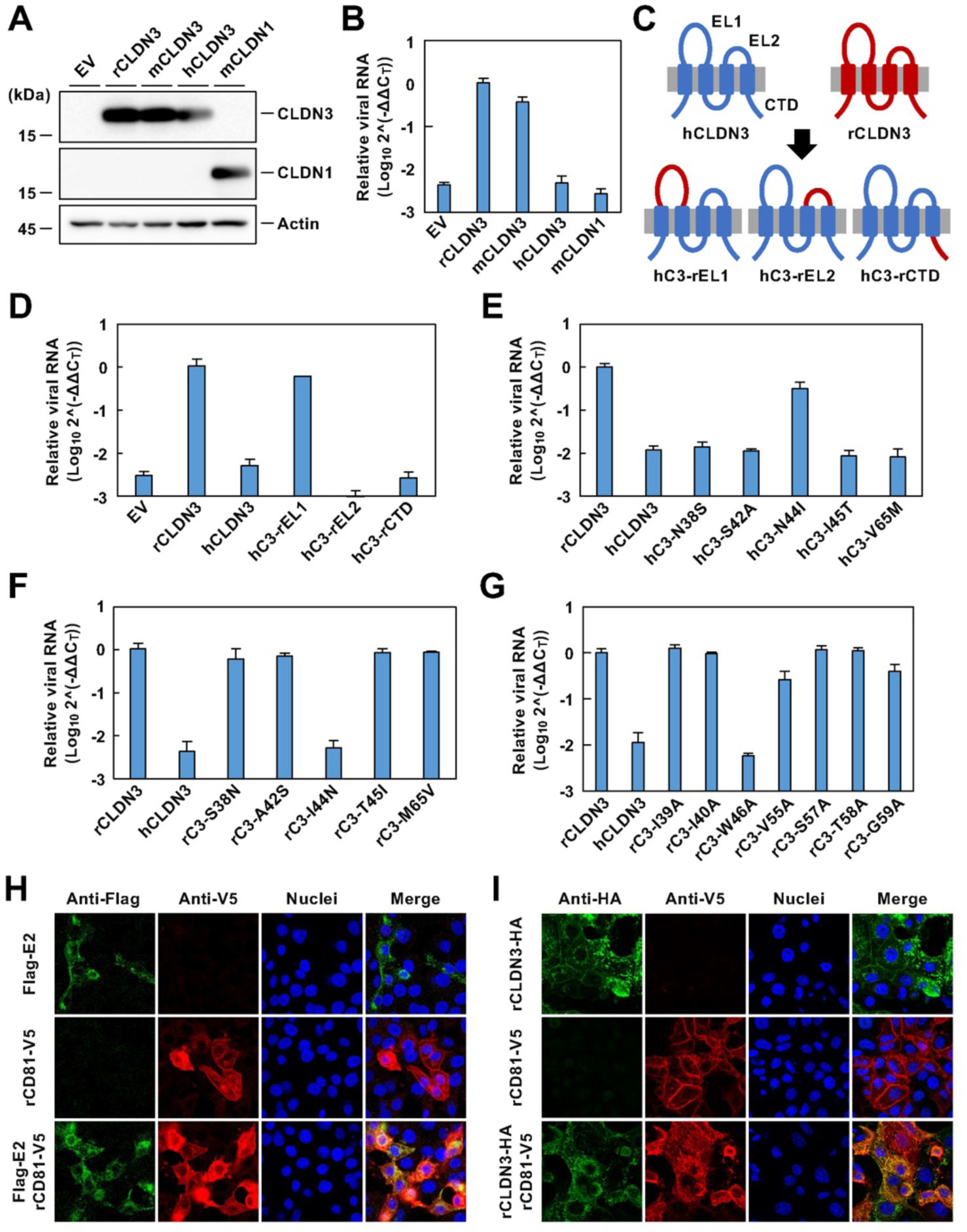
Identification of CLDN3 sites required to facilitate NRHV1 entry. (A) Western blot analysis of lysates extracted from AML/122 cells transfected with the indicated plasmids. (B) AML/122 cells were transfected with the indicated plasmid 24 h before inoculation with NRHV1cc at 5 copies/cell. Each transfected cell group was harvested at 3 dpi. Total RNA was subjected to qRT‒PCR to target the viral RNA. (C) Scheme of human-rat chimeric CLDN3 constructs. (D) AML/122 cells were transfected with the indicated plasmid 24 h before inoculation with NRHV1cc at 5 copies/cell. These cells were harvested at 3 dpi. Total RNA was subjected to qRT‒PCR to target the viral RNA. (E-G) AML/122 cells were transfected with plasmids encoding hCLDN3 mutants (E), rCLDN3 mutants (F), or rCLDN3 surface residue mutants (G) 24 h before inoculation with NRHV1cc at 5 copies/cell. These cells were harvested at 3 dpi and then subjected to qRT‒PCR to target the viral RNA. (H, I) AML12 cells were cotransfected with plasmids encoding V5-tagged rCD81 and Flag-tagged NRHV1 E2 genes or with plasmids encoding V5-tagged rCD81 and HA-tagged rCLDN3 genes. The cells were fixed at 24 h post-transfection and subjected to immunofluorescence analysis.

## Discussion

NRHV1 is known to infect immunocompetent rat strains, leading to chronic hepatitis, although other species have not yet been reported to be susceptible to infection in laboratory rat and murine strains ^8, 10^. Researchers believe that *in vivo* and *in vitro* infection models of NRHV1 will serve as useful surrogate viral models for HCV research. However, the virological characteristics of NRHV1 remain poorly understood. The NRHV1-permissible cell line from rat hepatoma McA-RH7777 cells was established recently, whereas the development of cells from mouse hepatocytes has not yet been attempted ^25, 26^. In this study, we also successfully established an NRHV1-permissible cell line, McA1.8, from the rat McA-RH7777 cell line harbouring NRHV1 SGR (Figs. 1, 2). An increase in NRHV1 replication was observed in McA1.8 cells, similar to reports regarding the increased efficacy of HCV replication in HCV replicon-cured cells ^35^, which may be attributed to the increased expression of miR-122 (Supplementary Figs. 1A, 2A and 2B). Infection of McA1.8 cells with NRHV1ser, which was collected from immunodeficient mice persistently infected with wild-type NRHV1, resulted in the production of supernatant NRHV1 RNA during approximately one month of serial passages (Fig. 2A). The infectious viral particles were included in the supernatant, since viral passage was possible (Fig. 2G, H). The resulting viral particle was cell culture adaptive and designated NRHV1cc in this study. The viral genome of NRHV1cc contains multiple nonsynonymous mutations (Supplementary Table S4). Mutations accumulated in the nonstructural gene region may contribute to the increased viral replication *in vitro*, although further studies are needed to confirm this phenomenon. The NRHV1cc strain prepared from the McA-RH7777.hi cell line included several adaptive mutations in NS3 and NS5A, such as S1757A/T2373A and K2130Q/E2223K/D2374G, that promoted the propagation of NRHV1cc ^26^, while these mutations were not observed in our NRHV1cc strain. In addition, the L586S mutation in E2 was common to both NRHV1cc strains, which may contribute to increasing viral infectivity in cultured cells.

Since murine Hepa1-6 cells support NRHV1 replication (Fig. 1) but do not permit NRHV1 propagation ^11^, AML12 cells were used in this study to establish a murine cell-derived culture model for NRHV1. Our data suggest that AML12 cells do not support NRHV1 replication and entry due to deficient expression of miR-122 and CLDN3, respectively (Figs. 1, 5). Furthermore, AML/122/rC3 cells supported NRHV1cc entry and replication but not the formation of infectious particles (Fig. 5), suggesting that the AML12 cell line lacks some factors required for the assembly of infectious viral particles. In contrast, previous reports have demonstrated that murine hepatoma cell lines such as AML12, MMH1-1, Hepa1-6 and Hep56.1D, but not rat McA-RH7777 cells, support low replication of HCV SGR without miR-122 replenishment ^45, 46^. The mechanism by which miR-122 facilitates hepacivirus replication has not been fully elucidated ^47^, and the dependency on miR-122 for viral replication seems to vary among hepacivirus species because NPHV replication is established using the stem loop I of the 5’UTR instead of miR-122 ^12, 13^. In the case of HCV, viral replication under miR-122 depletion or inactivation conditions induces the generation of a variant with a G28A mutation, which supports miR-122-independent replication, although miR-122 is required for sufficient HCV replication ^27, 47^. In addition, previous studies have shown that some HCV genotypes with adenine at position 28, such as genotypes 1b (Con1), 4a (ED43) and 5a (SA13), are resistant to miR-122 depletion in patients ^2, 27^. Our data suggest that the replication of the NRHV1 rn1 strain strongly depends on miR-122, unlike that of HCV and NPHV. Furthermore, murine AML12 and MMH1-1 cells do not secrete infectious JFH-1 particles following full-length JFH-1 RNA transfection ^45^. HCV assembly and release are associated with the very low-density lipoprotein (VLDL) biosynthesis pathway. AML12 cells expressing apolipoprotein E (apoE) can produce infectious particles in a trans-complementation assay employing AML12-derived SGR cells, suggesting that the lack of production of infectious HCV particles in AML12 cells is due to the loss of apoE ^48^. However, we cannot deny the possibility that factors other than apolipoprotein are required for infectious HCV particle formation in the AML12 cell line because compared with Huh7 cells, AML12 cells expressing apoE produce extremely low quantities of particles ^48^. The association of NRHV1 particle formation with VLDL synthesis has not yet been revealed and should be investigated in the future.

Recently, Wolfisberg and colleagues demonstrated that SR-BI is essential for NRHV1 entry ^25, 26^. The data in this study indicate that rodent CD81, OCLN and CLDN3, in addition to SR-BI, are utilized in the entry of NRHV1. The data deduced from single-cell RNA sequence analysis of the murine liver suggest that CLDN3, but not CLDN1, is a major member of the CLDN family in mouse hepatocytes ^36^. NRHV1 susceptibility was lost in McA1.8 cells deficient in OCLN and CLDN3 but was partially lost in CD81-knockout cells (Figs. 3, 4). These findings suggest that a host factor(s) compensates for the functions of CD81 under at least *in vitro* conditions. The expression of CLDN3, but not that of CLDN1, promoted the entry of NRHV1 into AML/122 cells (Fig. 5). Given that CLDN3 possesses structures homologous to those of CLDN1, NRHV1 may enter cells through a multistep process similar to HCV entry ^19^. Furthermore, the receptor function of CLDN3 corresponded with the host range of NRHV1 (Fig. 6B). In contrast, nonhost CLDN1 can facilitate HCV entry ^20^, although CLDN3 does not support HCV entry ^22, 23^. Although the role of CLDN1 in HCV entry is critical, our data indicate that CLDN3 plays a role in determining the host specificity of NRHV1. Determining which member of the CLDN family is predominant may contribute to the adaptive evolution of individual hepacivirus species. Interestingly, the AML/122 cell line, which is deficient in OCLN and CLDN3, allowed NRHV1 entry following CLDN3 replenishment alone and exhibited an increase in NRHV1 susceptibility by OCLN coexpression, but this increase was not notable (Fig. 5A, I). OCLN expression confers weak HCV susceptibility to AML12 cells ^20^. AML/122 cells may express an alternative factor or OCLN variant that facilitates NRHV1 entry. Given that human CD81 and OCLN are essential for HCV entry, further research is needed to elucidate the specific entry mechanisms of NRHV1.

The CLDN family comprises more than 20 members and shares a structural topology of four transmembrane and two extracellular loop domains. The entry step of NRHV1 was shown to be mediated by EL1 of CLDN3, similar to CLDN1, in the case of the HCV entry step (Fig. 6C), although GBV-B entry is suggested to be mediated by EL2 of CLDN1 ^24^. Complex formation of CD81 and CLDN1 through EL1 is indispensable for HCV entry ^44^. Since CD81 is dispensable for GBV-B entry ^24^, EL1 of CDLN3 may play an important role in the entry of NRHV1 through its interaction with CD81 or other equivalents. Our data indicated that Ile^44^, a specific residue for rat/mouse CLDN3, and Trp^46^, a conserved residue among rat/mouse and human CLDN3, are essential for NRHV1 entry (Fig. 6F, G). Trp^46^ of rat and murine CLDN3 is not included in the W^29-GLW50-C53-C63^ signature sequence conserved among EL1s of CLDN members (Supplementary Fig. 7A). The protein structure predicted by AlphaFold2 also underscores the importance of Ile^44^ and Trp^46^ in NRHV1 entry. According to the structure prediction, these residues are exposed on the molecular surface and are adjacent to each other. The amino acid substitution of Ile^44^ with Asn in murine CLDN3 likely alters the microstructure conformation involving Ile^44^ and Trp^46^ by generating an interaction between Asn^44^ and Phe^34^. Although the precise roles of CLDNs in the entry of hepaciviruses remain under investigation, previous findings on HCV entry suggest that CLDN1 functions at a postattachment step to facilitate endocytosis of the HCV-CD81 complex ^19, 44^. In fact, NRHV1 E2 and CD81, as well as CD81 and CLDN3, colocalized on the cell surface (Fig. 6H, I), suggesting that CLDN3 may facilitate the recruitment of the NRHV1-CD81 complex to the tight junction region, similar to the role of CLDN1 in HCV entry. Further analysis is needed to elucidate the involvement of the microstructure containing two adjacent amino acid residues, Ile^44^ and Trp^46^, in the postattachment step.

In conclusion, we established rat McA1.8 cells and mouse AML/122/rCLDN3 cells as cellular models for NRHV1 infection. McA1.8 cells can support the entire life cycle of NRHV1, while AML/122/rCLDN3 cells permit NRHV1 entry and replication but not particle formation. Our data also suggest that NRHV1 requires CLDN3 but not CLDN1 for entry in addition to SRB1, CD81 and OCLN in a partially similar way to HCV entry. CLDN3 may play a role in NRHV1 entry in cooperation with CD81. EL1 of CLDN3 is considered essential for NRHV1 entry. Our data regarding point mutations in CLDN3 suggest that the host-specific microstructure on the surface of CLDN3 is essential for CLDN3-mediated NRHV1 entry. Understanding virus‒host interactions will not only advance in the development of cell culture models but also provide insight into the host range, origin and evolution of hepacivirus.

## Acknowledgements

We would like to thank Amit Kapoor for providing the infectious clone of the RHV rn1 strain, Troels K. H. Scheel for providing SGR vectors, Naoki Ohtsu for technical assistance and helpful discussion, Masako Mori for secretarial work, and Midori Yoda and Keiko Ohmaki for technical assistance. This work was supported by the Japan Science and Technology Agency (JST) Moonshot R&D under grant number JPMJMS2025, by the Japan Agency for Medical Research and Development (AMED) under grant number JP24fk0210109, by JSPS KAKENHI Grant Number JP 15K18779, by the Takeda Science Foundation, and by a scholarship donation from Yakult Co., Ltd.

## Supplementary Information

### Supplementary Figure legends

**Supplementary Fig. 1.**
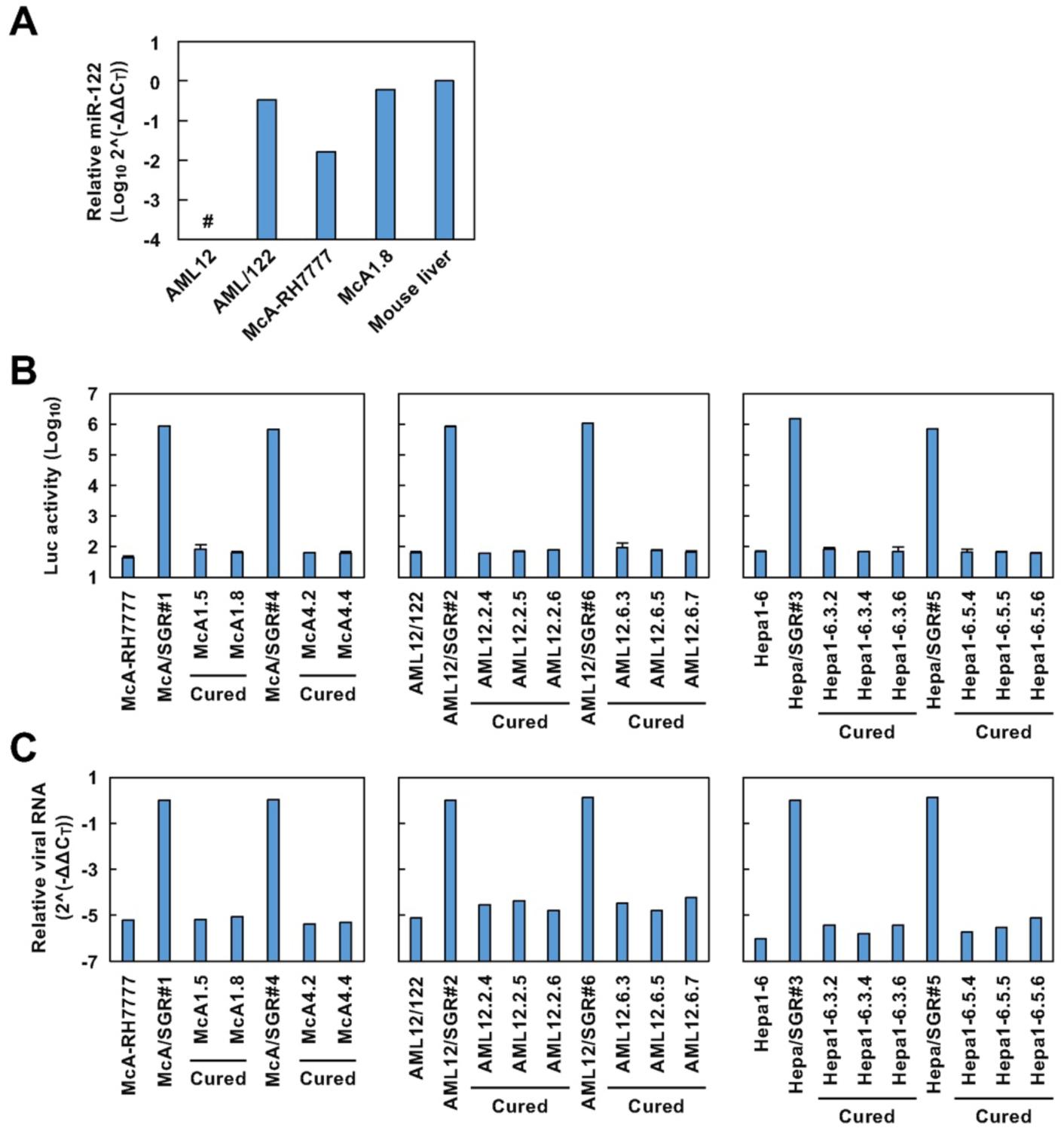
Expression levels of miR122 and NRHV1 SGR RNA in murine and rat cell lines. (A) The levels of miR122 relative to U6 RNA (ΔCt) were quantified via qRT‒PCR, normalized to the level in the mouse liver (ΔΔCt), and expressed as 2^−ΔΔCt^. Undetectable values are indicated by the number signs (#). (B, C) We established four or six cured cell clones from McA-RH7777, AML/122 and Hepa1-6-based replicon cells that harbour NRHV1 SGR. These cells were subjected to a luciferase assay (B) or qRT‒PCR targeting viral RNA (C). NRHV1 SGR RNA levels relative to those of GAPDH (ΔCt) were normalized to the levels in the cells harbouring NRHV1 SGR (ΔΔCt) and expressed as 2^−ΔΔCt^.

**Supplementary Fig. 2.**
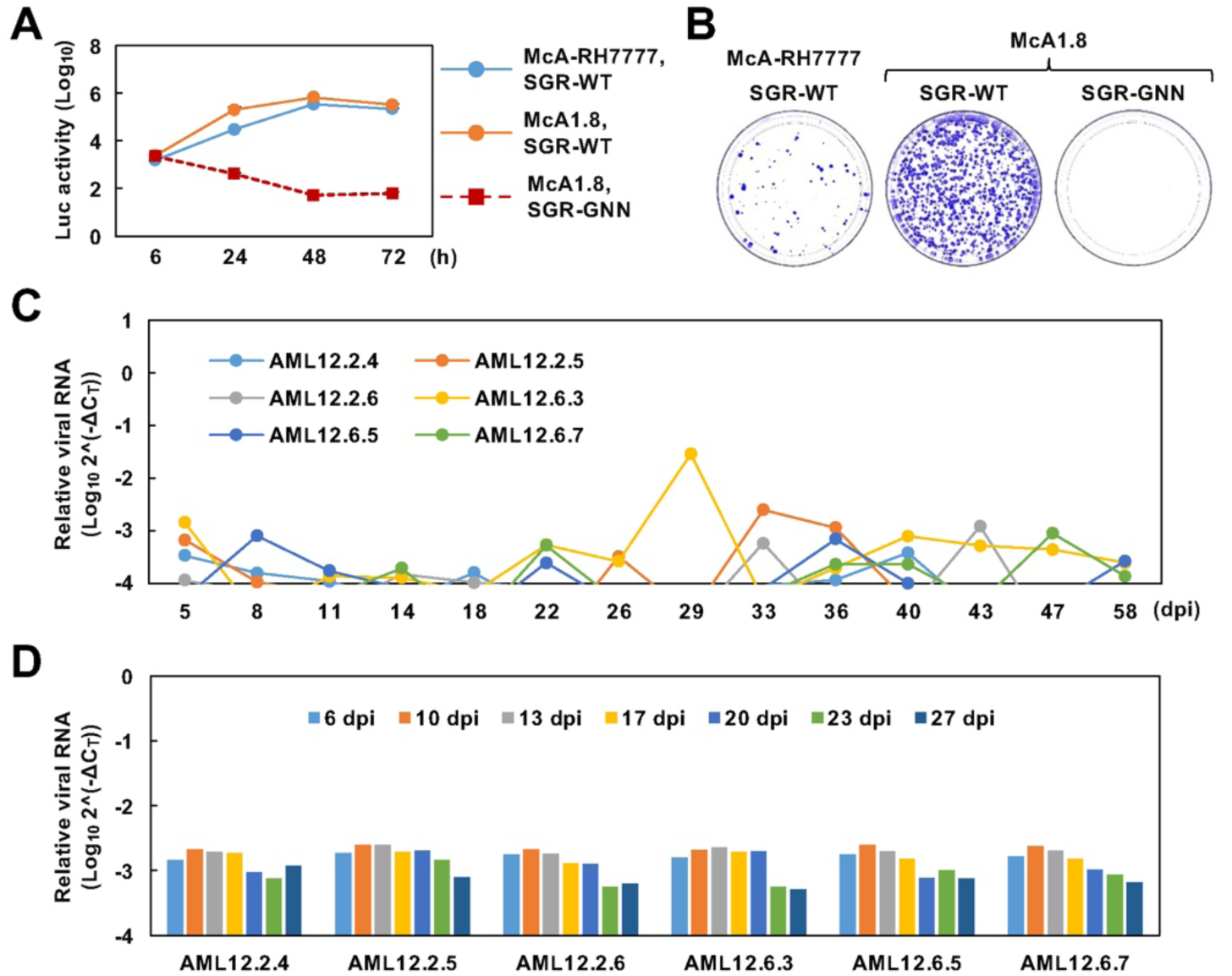
The permissibility of McA1.8 cells and AML/122-derived cured cell clones to NRHV1. (A, B) McA-RH7777 and McA1.8 cells were transfected with NRHV1 SGRs and then subjected to a luciferase assay at the indicated time points (A) or a colony formation assay (B). (C, D) Cured cell clones derived from AML/122 replicon cells were consecutively passaged every three or four days after inoculation with NRHV1ser at 6 copies/cell (C) or NRHV1cc prepared from McA1.8 cells at 10 copies/cell (D). The intracellular NRHV1 RNA level relative to the GAPDH mRNA level (ΔCt) was quantified via qRT‒PCR.

**Supplementary Fig. 3.**
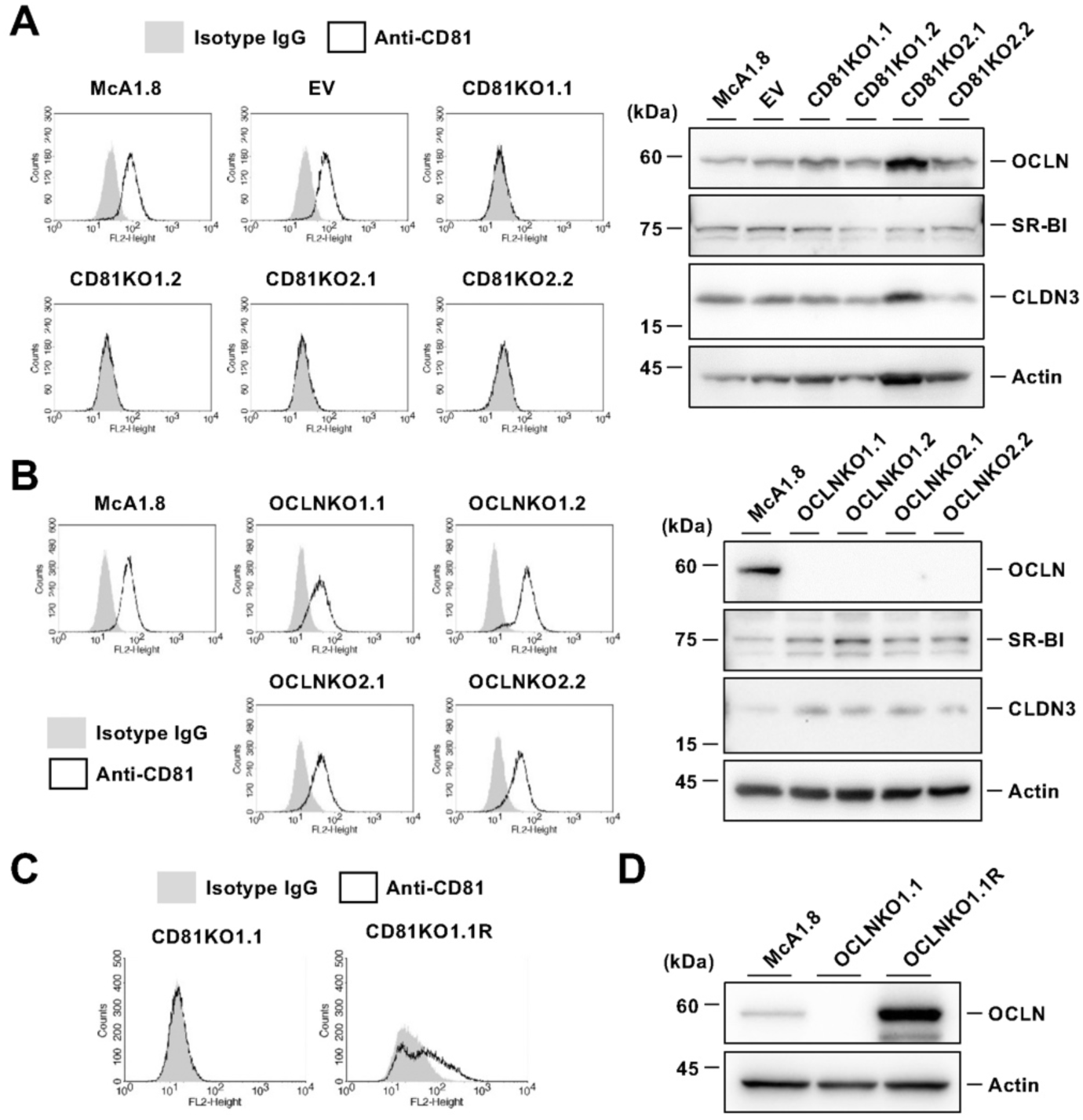
Establishment of CD81 and OCLN-KO cell lines from McA1.8 cells. (A) McA1.8 cells were transfected with plasmids expressing two different sgRNAs/Cas9s targeting the CD81 gene or the empty vector. The indicated cell clones were obtained following puromycin selection. Left panels: These cell clones were stained with PE-conjugated CD81-specific (black line) or isotype antibodies (grey shading) and analysed by flow cytometry. Right panels: Western blot analysis of lysates extracted from the indicated cell clones. (B) OCLN-deficient cell clones were obtained by the transfection of plasmids expressing two different sgRNAs/Cas9s targeting the OCLN gene and puromycin selection. Left panels: These cell clones were stained with PE-conjugated CD81-specific (black line) or isotype antibodies (grey) and analysed by flow cytometry. Right panels: Western blot analysis of lysates extracted from the indicated cell clones. (C) CD81KO1.1 cells were transduced with a retrovirus encoding sgRNA-resistant CD81 to obtain CD81KO1.1R cells. These cells were stained with PE-conjugated CD81-specific (black line) or isotype antibodies (grey) and analysed by flow cytometry. (D) OCLNKO1.1 cells were transduced with a retrovirus encoding sgRNA-resistant OCLN to obtain OCLNKO1.1R cells. These cell lysates were subjected to Western blot analysis.

**Supplementary Fig. 4.**
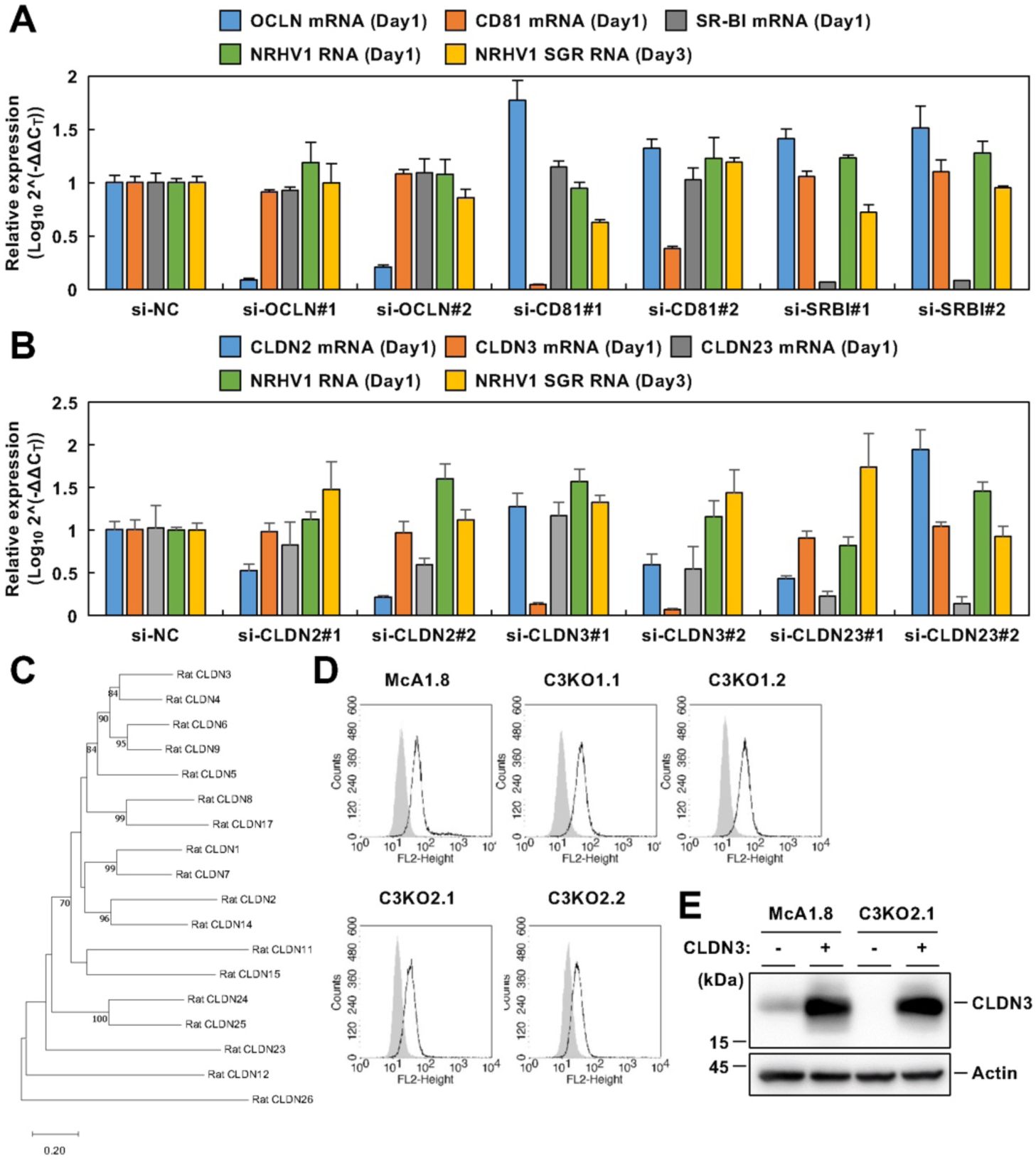
Knockdown, phylogenesis and expression of host factors that facilitate NRHV1 entry. (A, B) Effects of siRNA transfection on NRHV1 replication. McA/SGR#1 cells were transfected with 10 µM of each siRNA and harvested at 1 or 3 days post-transfection. The knockdown efficacy of each siRNA was evaluated at 1 day post-transfection. The level of NRHV1 SGR RNA was quantified at 1 and 3 days post-transfection. The relative RNA levels to the level of GAPDH mRNA (ΔCt) were normalized to the levels in cells transfected with si-NC (ΔΔCt). (C) A phylogenetic tree of amino acid sequences of rat CLDN members was generated by the neighbour‒joining method using MEGA11 software. Bootstrap values were calculated from 1000 replicates, and only bootstrap values of ≥70% are shown. (D) The indicated cell lines were stained with a PE-conjugated hamster anti-mouse/rat CD81 antibody (black line) or a PE-conjugated isotype IgG (grey) and analysed for CD81 expression by flow cytometry. (E) McA1.8 and C3KO1.1 cells were transfected with a plasmid encoding sgRNA-resistant CLDN3 (+) or an empty vector (-) and then subjected to Western blot analysis.

**Supplementary Fig. 5.**
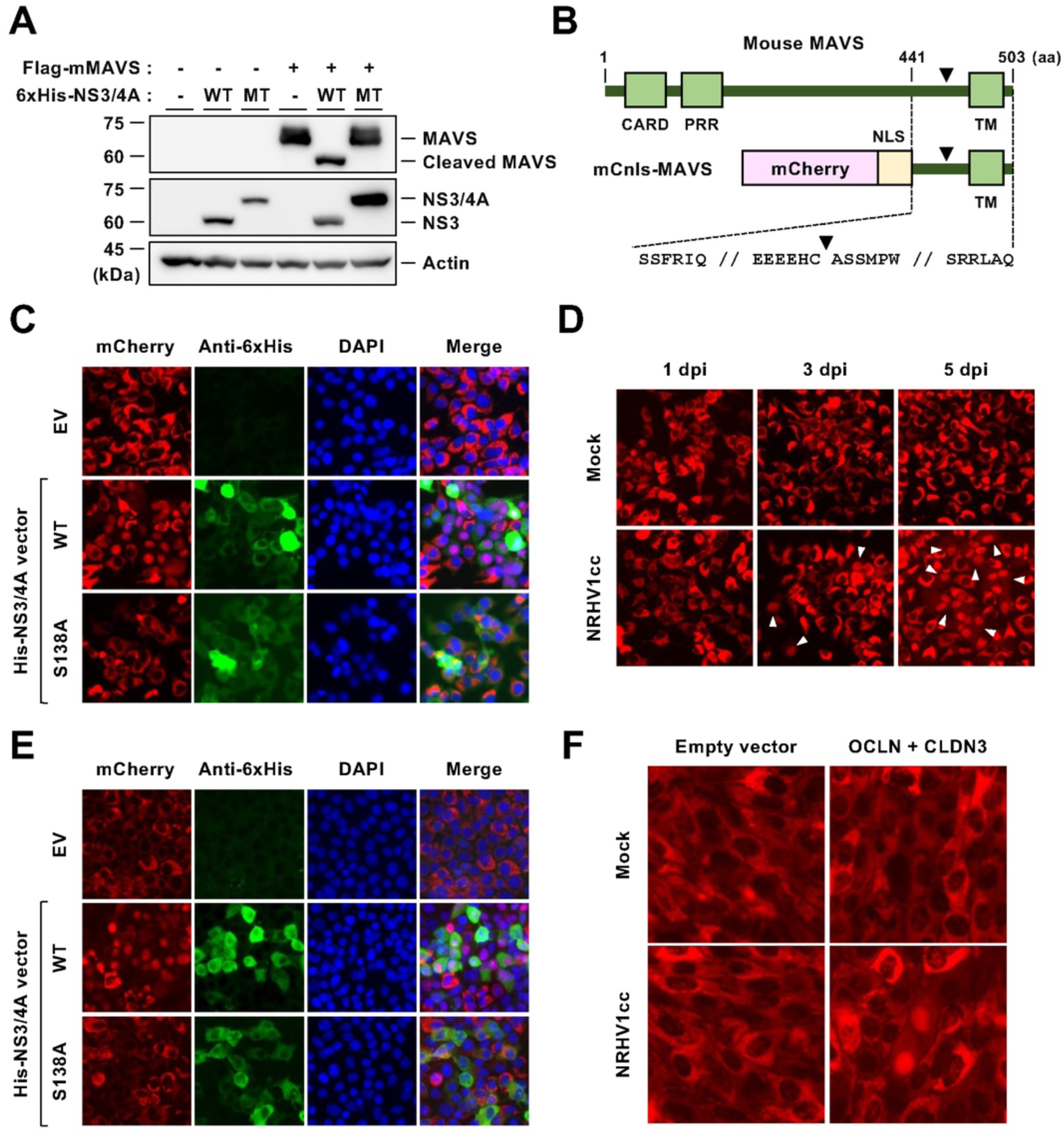
A live imaging system to detect NRHV1-infected cells. (A) McA1.8 cells were cotransfected with a plasmid encoding N-terminally Flag-tagged mouse MAVS (Flag-mMAVS) and a plasmid encoding N-terminally 6 × His-tagged NRHV1 NS3/4A (WT) or the S138A mutant (MT) in the indicated combinations. These cells were harvested at 24 h post-transfection and subjected to Western blot analysis. (B) Schematic diagrams of mouse MAVS (top panel) and the fluorescent reporter protein mCnls-MAVS (middle panel), which is composed of mCherry, SV40 NLS and MAVS. The bottom letters show the amino acid (aa) sequence of the region comprising aa 441-503 of mouse MAVS. The cleavage site of MAVS by NS3/4A is indicated by arrowheads. CARD, caspase recruitment domain. PRR, proline-rich region. TM, transmembrane domain. (C) McA1.8 cells stably expressing mCnls-MAVS (McA1.8/mCnls-MAVS) were transfected with plasmids encoding WT or S138A NS3/4A. These cells were fixed at 24 h post-transfection and stained with an anti-6 × His-tag antibody followed by an Alexa Fluor 488-conjugated anti-mouse IgG antibody (green). Nuclei were stained with DAPI (blue). (D) McA1.8/mCnls-MAVS cells were incubated with NRHV1cc, and the medium was subsequently replaced 24 h after infection. Images were captured at the indicated time points. Arrow heads indicate the nuclear translocation of mCnls. (E) AML/122 cells stably expressing mCnls-MAVS (AML/122/mCnls-MAVS) were transfected with plasmids encoding WT or S138A NS3/4A. These cells were fixed at 24 h post-transfection and stained in the same way as in (C). (F) AML/122/mCnls-MAVS cells were cotransfected with plasmids encoding rat OCLN and CLDN3 or the empty vector for 24 h. Then, these cells were inoculated with NRHV1cc, and the medium was replaced 24 h after infection. Images were captured at 4 dpi.

**Supplementary Fig. 6.**
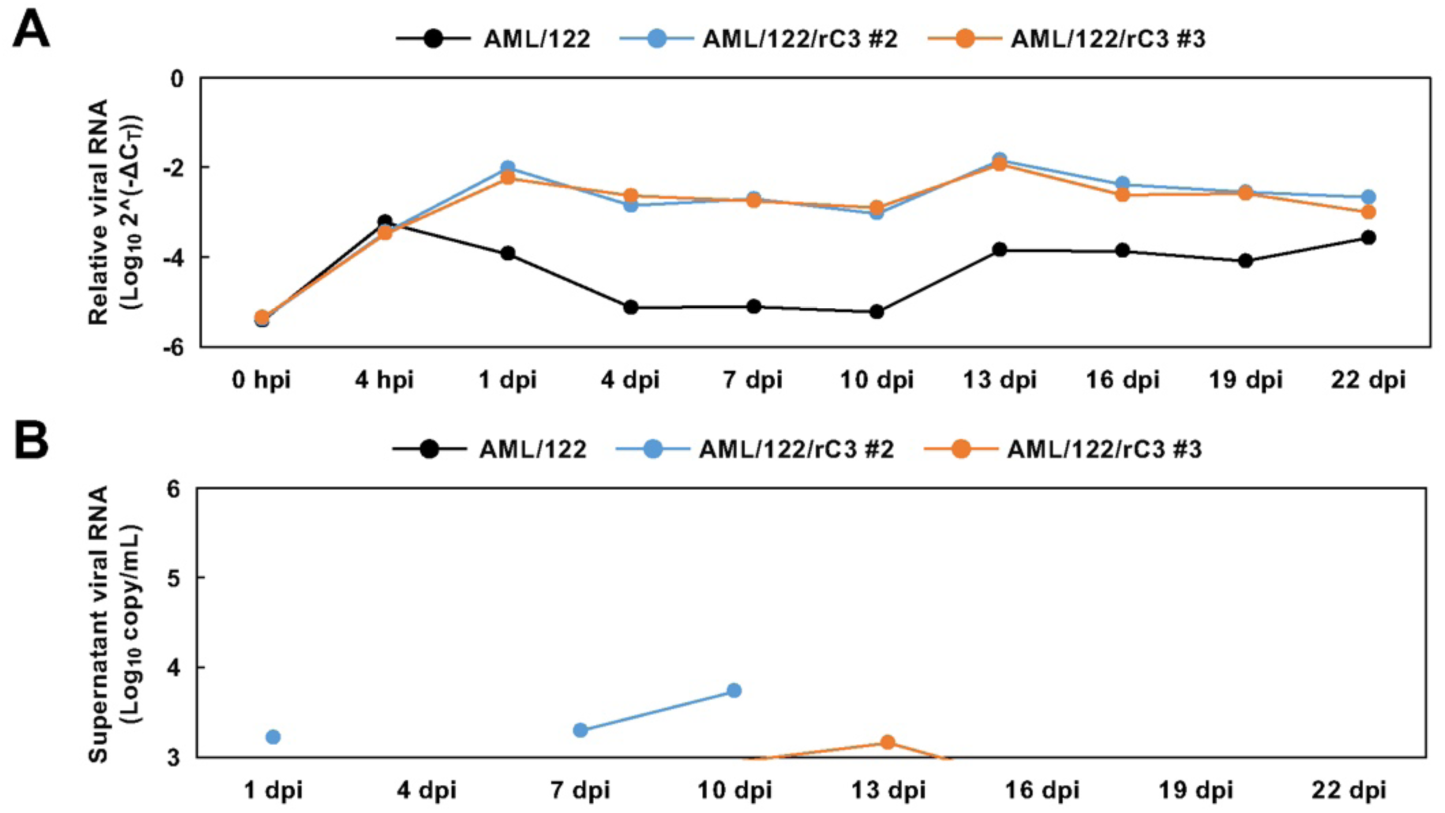
Time course of NRHV1 RNA level in AML12-derived cells. The cells were infected with NRHV1cc at 10 copies/cell for 4 h and washed with PBS to remove residual virus. These cells were then passaged every three or four days. At the indicated time points, NRHV1 RNA in the cells (A) and supernatants (B) was quantified via qRT‒PCR. The levels of intracellular NRHV1 RNA were plotted as values relative to the level of GAPDH mRNA (ΔCt).

**Supplementary Fig. 7.**
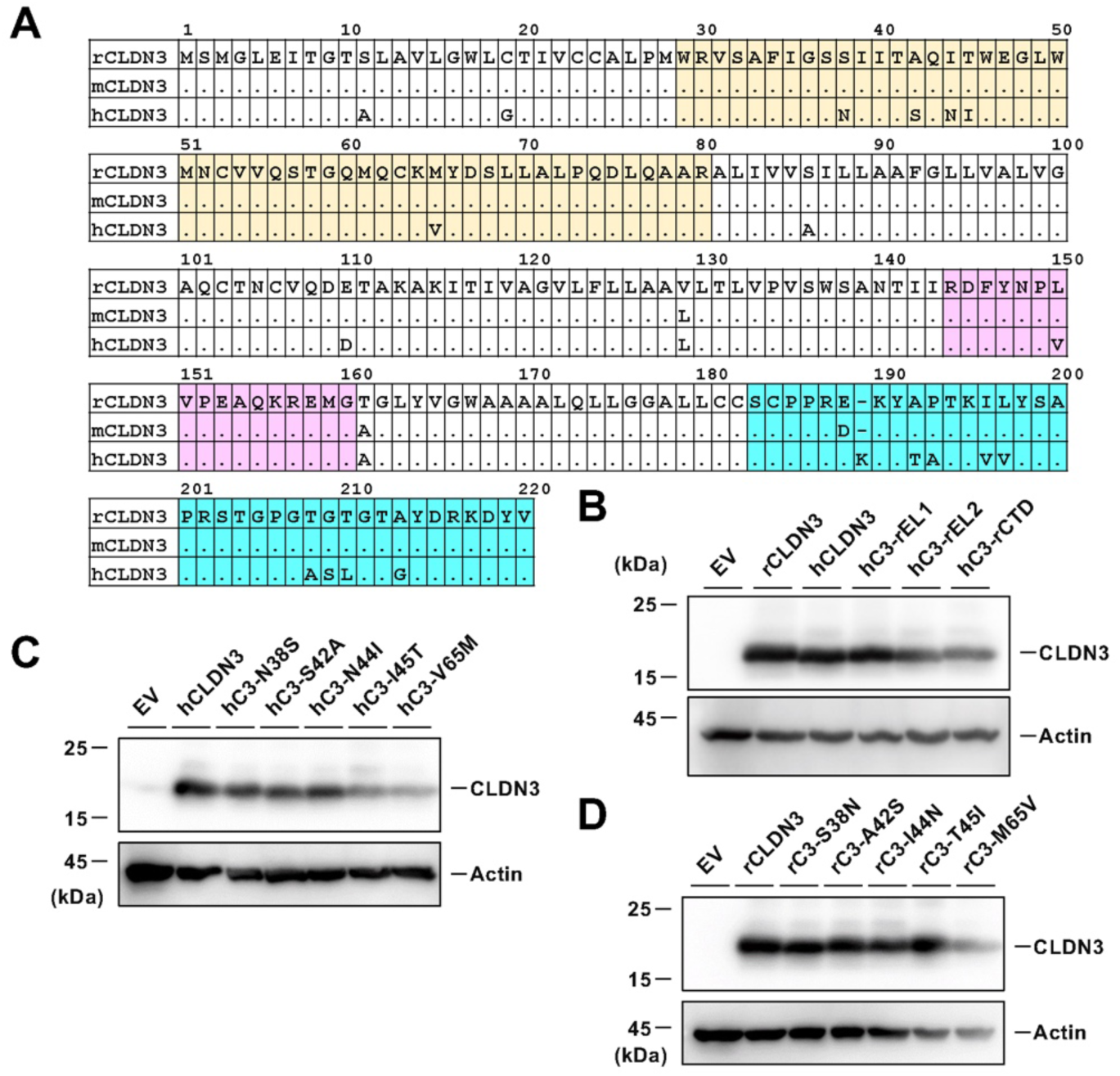
Amino acid sequences of CLDN3 and expression of CLDN3 mutants. (A) Amino acid sequences of rCLDN3, mCLDN3 and hCLDN3 are aligned. EL1, EL2 and the C-terminal domain (CTD) are indicated by yellow, red and blue boxes, respectively. (B-D) AML/122 cells were transfected with the indicated plasmids. The cells were harvested at 24 h post-transfection and subjected to Western blot analysis.

**Supplementary Fig. 8.**
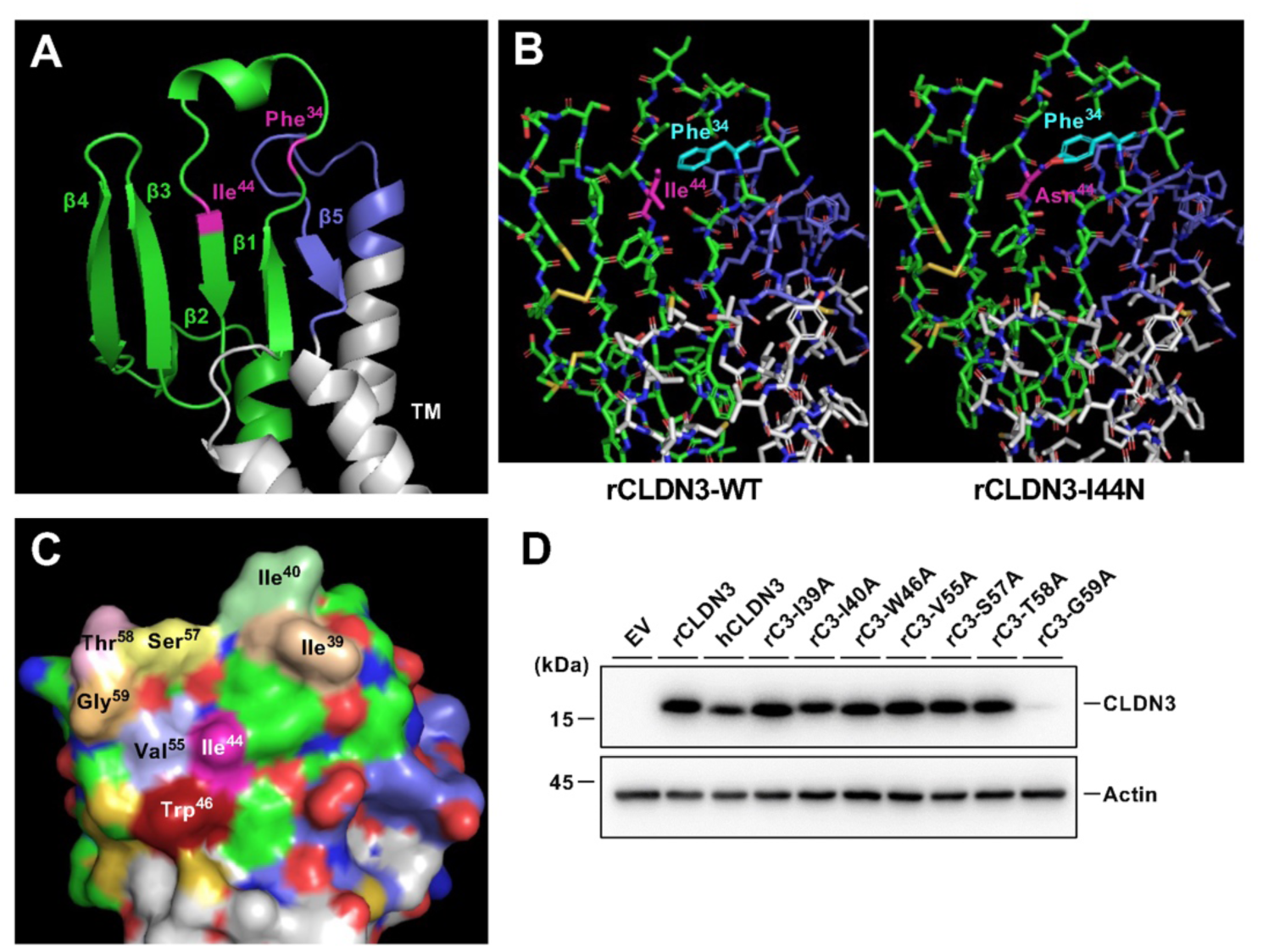
Amino acid sequences of CLDN3 and expression of CLDN3 mutants. (A) AlphaFold2 prediction of rCLDN3 shown in the cartoon model. EL1 and EL2 are coloured green and blue, respectively. Ile^44^ and Phe^34^ are highlighted in magenta. TM, transmembrane domain (grey). (B) AlphaFold2 prediction of wild-type rCLDN3 (left panel) and the I44N mutant (right panel) shown in the stick model. rCLDN3. Ile^44^ and Phe^34^ are displayed in magenta and cyan, respectively. Green, blue and grey represent EL1, EL2 and TM, respectively. (C) Key amino acid residues that compose the molecular surface of EL1 are indicated in the predicted surface model. (D) AML/122 cells were transfected with the indicated plasmids. These cells were harvested at 24 h post-transfection and subjected to Western blot analysis.

**Supplementary Table S1.**
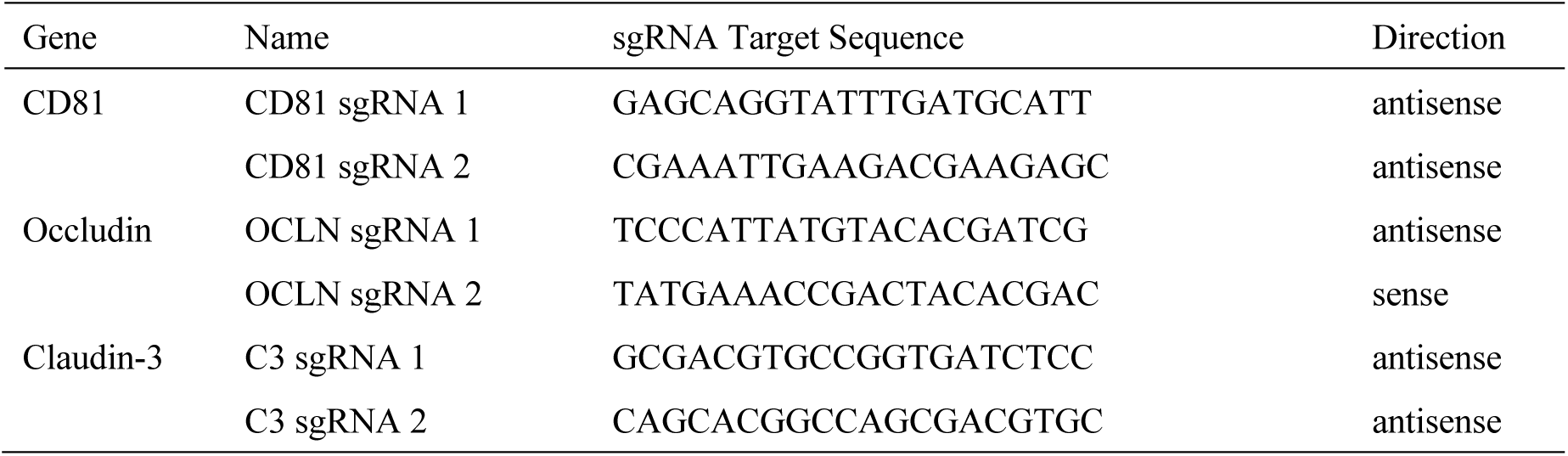
Synthesized sgRNA oligonucleotides for CRIPSR/Cas9 technology.

**Supplementary Table S2.** List of siRNAs.

| Name | Catalog No | Targeted gene |
| --- | --- | --- |
| si-CD81#1 | SASI_Rn01_00066147 | NM_013087 |
| si-CD81#2 | SASI_Rn01_00066149 |  |
| si-OCLN#1 | SASI_Rn01_00096936 | NM_031329 |
| si-OCLN#2 | SASI_Rn01_00096938 |  |
| si-SRBI#1 | SASI_Rn01_00079396 | NM_031541 |
| si-SRBI#2 | SASI_Rn01_00079397 |  |
| si-CLDN2#1 | SASI_Rn02_00222976 | NM_001106846 |
| si-CLDN2#2 | SASI_Rn02_00222977 |  |
| si-CLDN3#1 | SASI_Rn01_00104958 | NM_031700 |
| si-CLDN3#2 | SASI_Rn02_00265055 |  |
| si-CLDN23#1 | SASI_Rn01_00076732 | NM_001033062 |
| si-CLDN23#2 | SASI_Rn01_00076733 |  |

**Supplementary Table S3.**
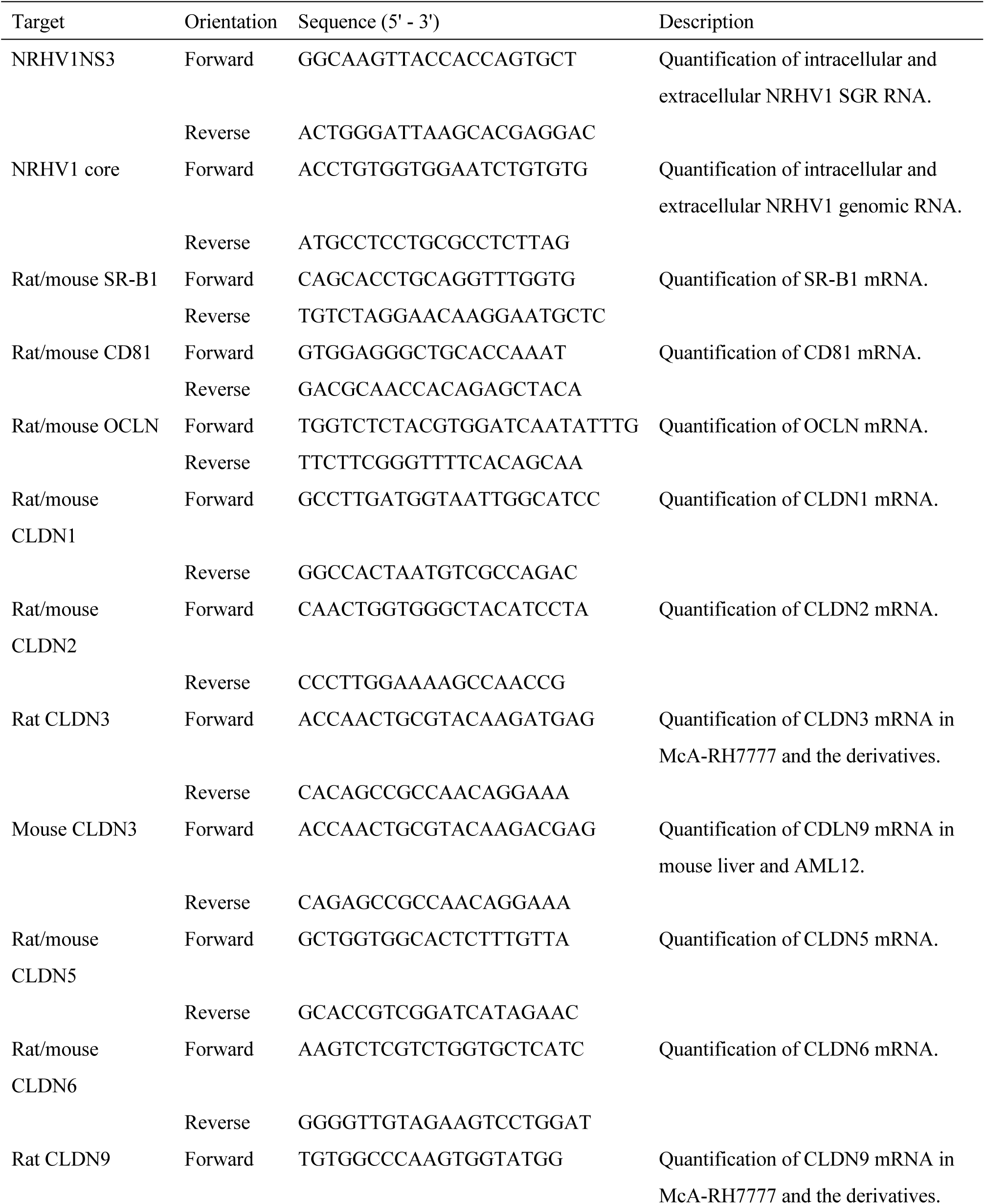
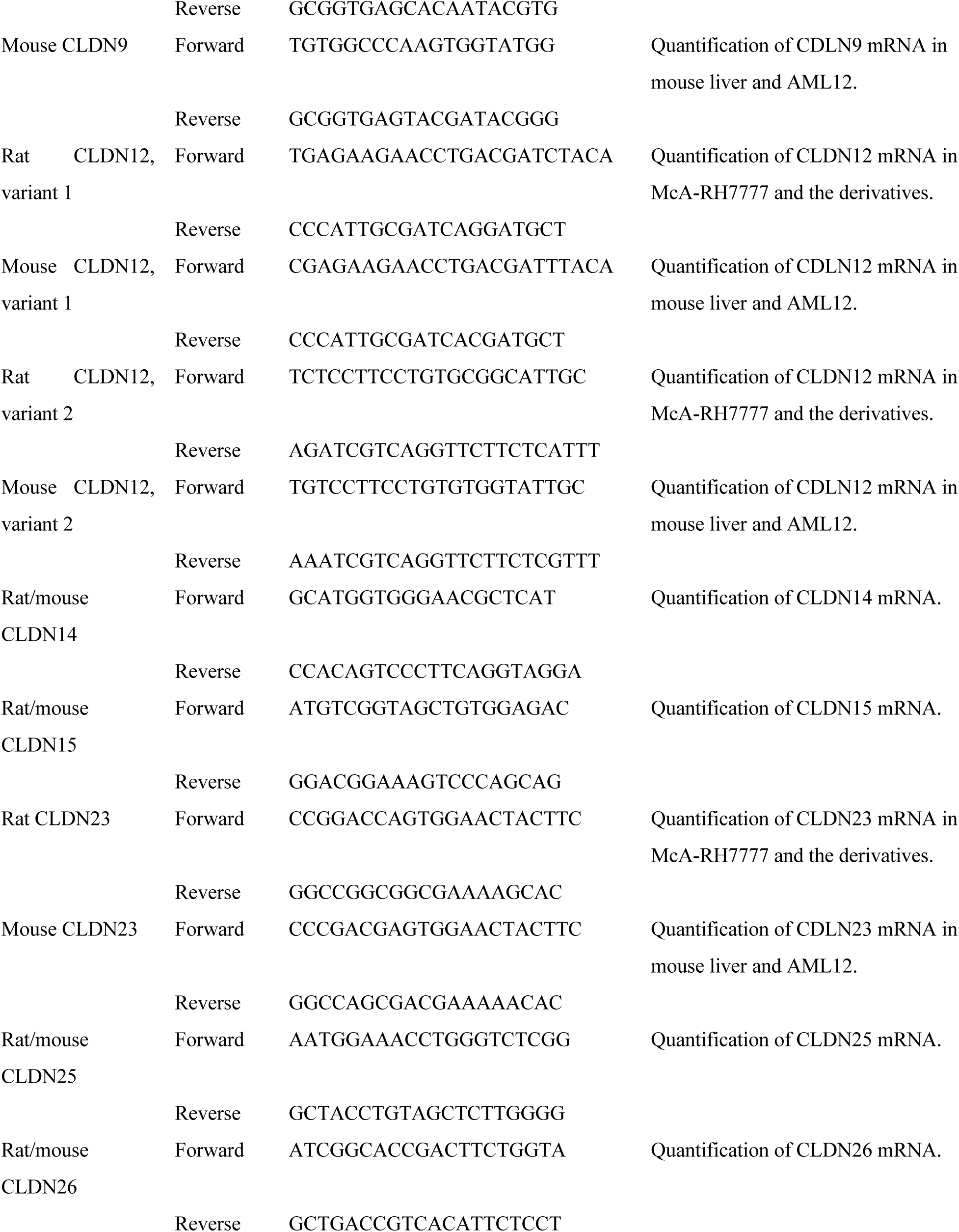
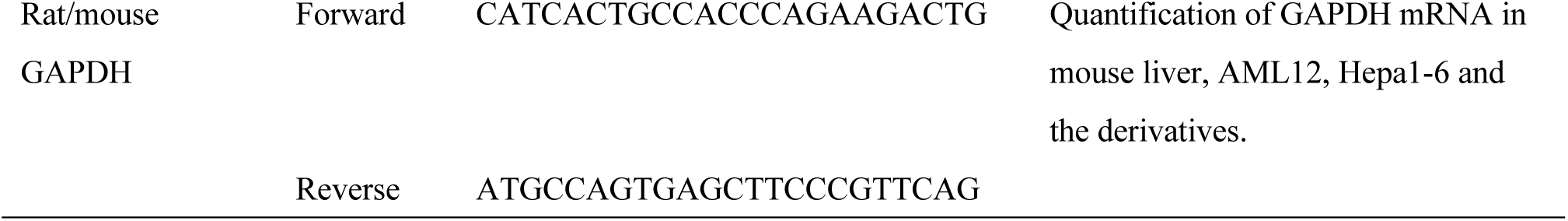
Primers for qRT-PCR.

**Supplementary Table S4.** List of synonymous and nonsynonymous mutations observed in NRHV1cc.

| Region | Changes in nucleotides |  |  | Changes in amino acids |  |  | Frequency (%) |
| --- | --- | --- | --- | --- | --- | --- | --- |
|  | Position | Inoculum | Change | Position | Inoculum | Change |  |
| 5' UTR | 79 | T | Del <sup>1)</sup> |  |  |  | 97.13 |
| E2 | 1781 | G | A | 432 | Q | - <sup>2)</sup> | 95.91 |
|  | 1849 | T | C | 455 | L | P | 80.32 |
|  | 2242 | T | C | 586 | L | S | 99.58 |
|  | 2321 | A | G | 612 | S | - | 99.27 |
| NS3 | 3664 | A | G | 1060 | K | R | 97.95 |
|  | 3878 | A | G | 1131 | L | - | 99.09 |
|  | 4082 | T | C | 1199 | T | - | 97.28 |
|  | 4287 | A | G | 1267 | T | A | 99.12 |
|  | 4637 | C | T | 1384 | F | - | 98.38 |
| NS5A | 6375 | G | A | 1963 | V | I | 98.51 |
|  | 7121 | T | C | 2212 | V | - | 96.26 |
|  | 7135 | T | C | 2217 | F | S | 97.5 |
|  | 7362 | T | C | 2292 | S | P | 88.6 |
|  | 7459 | A | G | 2325 | Q | R | 98.79 |
|  | 7486 | A | G | 2334 | K | R | 88.87 |
| NS5B | 7661 | A | C | 2392 | Q | H | 98.82 |
|  | 7862 | A | G | 2459 | T | - | 98.86 |
|  | 9022 | C | T | 2846 | A | V | 98.94 |

1) Single nucleotide deletion
2) Synonymoous mutation

